# Discovery and Engineering of a Therapeutic Interfering Particle (TIP): a combination self-renewing antiviral

**DOI:** 10.1101/820456

**Authors:** Elizabeth J Tanner, Seung-Yong Jung, Joshua Glazier, Cassandra Thompson, Yuqi Zhou, Benjamin Martin, Hye-In Son, James L Riley, Leor S Weinberger

## Abstract

Population-level control of HIV-1 faces recognized challenges, including the evolution of viral resistance and adherence issues in resource-limited settings. It has long been proposed that viral deletion mutants that conditionally self-renew at the expense of the wild-type virus (i.e., Defective Interfering Particles, DIPs^1^) could constitute a long-term intervention that circumvents adherence challenges and has a high genetic barrier to resistance, echoing recent approaches^2^. Theories predict^3, 4^ that DIPs could be engineered into a therapy for HIV-1 (i.e., *Therapeutic* interfering particles or ‘TIPs’) provided they stably persist in patients (R_0_>1) by spreading to new cells during active infection (hence, a self-replenishing antiviral). To date, DIPs amenable to such engineering have remained elusive for HIV-1. Here we report the discovery of an HIV-1 DIP and its subsequent engineering into a TIP. The TIP interferes with HIV-1 replication at multiple stages of the viral lifecycle, including genome packaging, virion maturation, and reverse transcription, essentially acting as a combination antiviral. In humanized mice, the TIP suppressed HIV-1 replication by ten-fold and significantly protected CD4+ T cells from HIV-1 mediated depletion. These data provide proof-of-concept for a class of biologic with the potential to circumvent significant barriers to HIV-1 control.

---

Despite the availability of effective antiretroviral therapy regimens, 37 million people are currently living with HIV-1/AIDS (hereafter HIV) and ∼2 million new HIV infections occur each year, largely in the developing world^5^. Epidemiological models^6^ project that epidemic control in sub-Saharan Africa using conventional approaches may be exceptionally challenging, due in part to deployment, behavioral, and mutational barriers^7–9^. It has been proposed that Defective Interfering Particles (DIPs) could overcome these barriers^3, 4, 10, 11^. Originally observed for influenza virus in the 1940s by von Magnus^12^, DIPs are viral deletion mutants that conditionally replicate (i.e., lack self-replication) and act as molecular parasites of the wild-type virus within infected cells, stealing essential proteins from wild-type virus (e.g., replication or packaging proteins). Consequently, DIPs suppress wild-type viral burst size (the number of viral particles released from an infected cell) and conditionally mobilize their own genomes, spreading their antiviral properties to new cells. Mathematical models predicted that DIPs could be long-acting therapeutics against HIV provided they can be synthetically engineered to that have a basic reproductive ratio [R_0_]>1, which have been called *Therapeutic* Interfering Particles (TIPs)^3, 4^ (Fig. 1a,b). R_0_ is a threshold value defined as the expected number of secondary infections arising from a single infected individual or cell^13^; a spreading infection has an R_0_>1 while the infection dies out at R_0_<1. The R_0_ has been used to define the threshold for TIP persistence; the TIP therapy will ‘spread’ through the host-cell population when R_0_^TIP^>1^3, 4, 11^. Previous work identified retroviral vectors with the potential to suppress HIV replication^14–18^, but a number of shortcomings, including suboptimal persistence, limited their translation into therapy^15, 19^.

**Figure 1:**
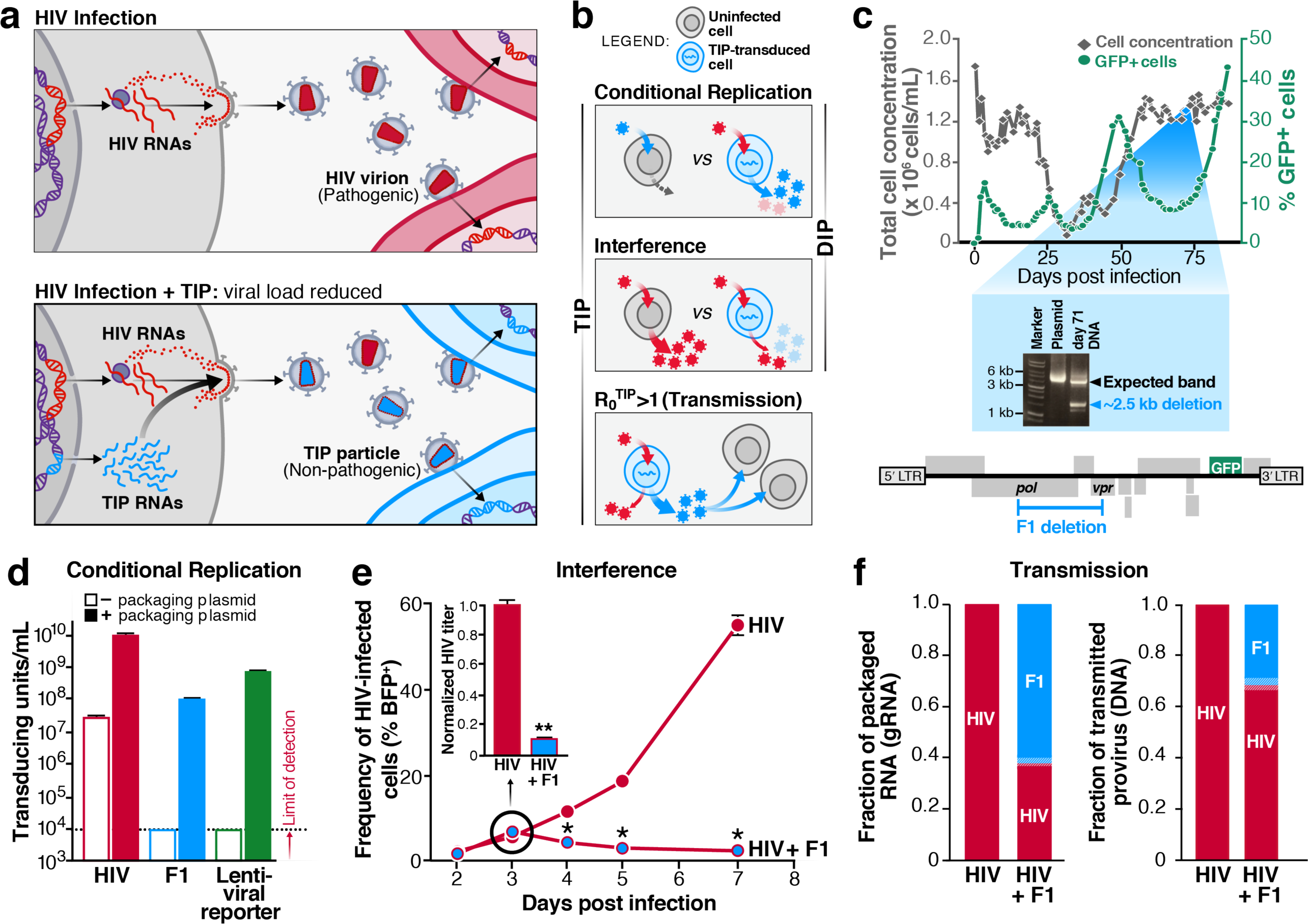
Long-term selection of an HIV defective interfering particle (DIP). **a**, Schematic of the TIP concept: replication-defective TIP RNAs hijack HIV packaging components and spread to new cells at the expense of the wild-type virus. TIP virus-like particles (VLPs) are integration competent but not toxic to host cells. **b**, Schematic of DIP vs. TIP criteria: (i) Conditional replication: DIPs and TIPs only replicate in the presence of HIV superinfection; (ii) Interference: DIPs and TIPs reduce HIV burst size; (iii) Transmission: TIPs must mobilize to more than one naïve cell (R_0_>1) in the presence of HIV. **c**, Detection of an HIV deletion variant in long-term culture. Top: cell concentration and frequency of GFP^+^ cells in long-term reactor culture. CEM CD4+ cells (hereafter “CEMs”) were maintained at steady-state growth in a reactor by daily replacement of approximately 25% of the total culture volume with fresh media (Supplementary Fig. 1b) prior to infection with HIV (day - 14 to day 0, not shown). On day 0, reactor was infected with HIV-GFP – molecular clone NL4-3 encoding a GFP reporter in the *nef* reading frame – at low MOI and cells were collected for analysis at regular intervals. Middle: PCR products obtained from integrated HIV proviral DNA. Total DNA was extracted from reactor cells on day 71 and analyzed with primers to the HIV proviral sequence (lane 3). Both the expected band (see HIV plasmid PCR, lane 2) and a band approximately 2.5 kb smaller are indicated on the gel image, which has been cropped to show relevant lanes and bands. Bottom: the “F1 deletion” mapped onto the HIV parent backbone. The location of the deletion was identified by submitting the small PCR band for Sanger sequencing. **d**, Transduction competence of F1 VLPs assembled in the presence and absence of packaging plasmid. The F1 deletion was cloned into the HIV-GFP backbone [construct name: F1(+Tat)], and the plasmid was transfected into 293T cells in the absence or presence of the Δ8.91R packaging plasmid and pseudotyped with VSVG to generate VLPs. VLPs were titered on permissive MT-4 cells by flow cytometry for GFP. Replication-competent HIV-GFP and a lentiviral reporter vector that expresses GFP from the EF1A promoter (pLEG, see table of constructs) were included as controls. **e**, Outgrowth of HIV in the presence and absence of the F1 construct. A F1-GFP cell line, transduced with single integrations of F1-GFP (at polyclonal loci), or a mock-transduced control, were infected with HIV-BFP at low MOI. The frequency of BFP^+^ cells was measured by flow cytometry at indicated timepoints. Inset: Filtered supernatants, harvested at 3 dpi, were titered by flow cytometry (48 after infection of naïve cells). Main panel: *p = 0.004 (4 dpi), p = 0.001 (5 dpi), p = 0.001 (7 dpi); Inset: **p = 5 x 10^-6^. All p values obtained using a two-tailed, unpaired Student’s t test. **f**, The fraction of F1 and HIV packaged genomic RNA in filtered supernatant (left) compared to the fraction of transmitted proviral DNA that integrated into new cells in a second round of infection (right). gRNA in the supernatant was analyzed by allele-specific RT-qPCR from samples panel e (inset). The transmitted DNA was measured by infecting MT-4 cells as if to titer (as in e), harvesting gDNA at 2 dpi (to allow for integration of WT and F1 constructs) and quantifying by allele-specific qPCR. All data represent the mean and s.e.m. of three biological replicates except in panel c. For long-term reactor culture data in panel c only one reactor trajectory is shown because trajectories and oscillations varied widely between reactors; trajectories for parallel reactors available on request.

To understand why mobilizing DIPs might be rare for HIV—unlike other RNA viruses where DIPs are readily isolated in high-MOI passage^12, 20^—we used established *in silico* models of HIV *in vivo* dynamics^21, 22^. The model predicted that a DIP spontaneously generated on day one would require 50–100 days of continuous culture to reach detectable levels (Supplementary Fig. 1a and Supplementary Methods) and would fail to expand in conventional serial-passage cultures where DIP-integrated cells would be diluted by exogenous replacement of new target cells. Based on these predictions, we designed an *in vitro* continuous-culture reactor (Supplementary Fig. 1b,c and Supplementary Methods) where CD4+ target cells are replenished exclusively by cell division. This reactor enabled long-term HIV culturing and allowed putative DIP-integrated cells to expand without dilution from exogenous replacement of cells.

The reactor was infected with an HIV GFP reporter virus and over the course of ∼90 days HIV dynamics, tracked by viral GFP expression, exhibited von Magnus-like fluctuations— stereotyped waves of viral amplification and collapse indicative of DIP formation^23^ (Fig. 1c, top). Integrated HIV proviral DNA from a late time point (day 71) was analyzed by PCR and revealed that a nearly 2.5kb region had been deleted from the viral genome (Fig. 1c, middle and bottom) and sequencing mapped this deletion—designated the “F1” deletion (for reactor “Flask #1”)—to the *pol–vpr* region of HIV. A similar deletion arose in a parallel reactor and sequence analysis showed that these deletions were flanked by short homology regions, indicating a likely recombination event (Supplementary Fig. 1d).

To test if the F1 deletion variant satisfied the requirements for a DIP (Fig. 1b), the F1 deletion was cloned into a GFP-tagged HIV backbone (Supplementary Fig. 2a), hereafter termed “F1”, and its interaction with HIV characterized in cell culture.

First, to test if F1 satisfied the conditional-replication requirement, we provided the deleted HIV-1 proteins to F1 by co-transfection with a packaging plasmid and quantified transduction-competent viral-like particles (VLPs). As expected, F1 only generated transduction-competent VLPs when the deleted HIV-1 proteins were provided in *trans* (Fig. 1d).

Second, to test F1’s ability to interfere with HIV, we used established approaches^24^ to generate an F1-transduced cell line—in which F1 RNA is expressed from single, polyclonal integration sites and is transactivated by HIV Tat protein delivered *in trans* by HIV superinfection (Supplementary Fig. 2a,b)—and measured the outgrowth of BFP-tagged HIV from these cells (for gating scheme see Supplementary Fig. 2c). Cells with and without F1 appeared equally permissive for initial HIV infection (see Fig. 1e [2 and 3 days post infection] and Supplementary Fig. 2d), but the subsequent outgrowth of HIV was restricted in the F1-integrated cell line, suggesting that F1 interfered with the spread of HIV. Titers of single-round infections also supported F1 interference with HIV; F1 reduced HIV titers by 94% in a single round of infection (p = 5 x 10^-6^) (Fig. 1e, inset) whereas no interference was observed in cell lines transduced with lentiviral vector controls (Supplementary Fig. 3a,b). To be sure that F1-associated interference was not cell-line dependent, we tested F1 interference in a more permissive T-cell line, MT-4 cells (Supplementary Fig, 3c), as well as human primary CD4+ T cells (Supplementary Fig. 3d) and found F1 significantly reduced HIV outgrowth in both settings.

Finally, to test if F1 RNAs mobilize in the presence of HIV (e.g., via stealing of packaging proteins from full-length HIV), we analyzed the packaged RNA from single-round infections (titered in Fig. 1e) by allele-specific qPCR. F1 RNA was present in extracellular VLPs and represented 60% of the total RNA population, showing a packaging advantage over HIV genomes (Fig. 1f, left).

Overall, these data verify that the F1 construct conditionally replicates (only in the presence of HIV), interferes with HIV outgrowth, and mobilizes from infected cells—thereby constituting a bona-fide DIP. However, despite F1’s packaging advantage, F1 exhibited a transmission defect, constituting only 30% of integrated provirus in the subsequent round of infection (Fig. 1f, right). Since TIP persistence and efficacy depend upon high R_0_^TIP^, we sought to improve transmission efficiency of the F1 construct.

In an effort to rationally enhance F1 transmission efficiency, we next examined F1’s genetic sequence. Given the location of the F1 deletion, which spans HIV’s central polypurine tract (cPPT), we hypothesized that the loss in transmission efficiency could be due to a reverse transcription or integration defect, since the cPPT has been shown to enhance lentiviral vector transduction efficiency^25–29^ and a recent forward genetic screen argues that the cPPT may be even more critical for viral transmission than previously appreciated^30^. Thus, we re-introduced the cPPT back into the F1 construct (hereafter termed “F1^cPPT^”; Fig 2a). F1^cPPT^ generated a two-fold increase in VLP-transduction efficiency relative to F1 (Fig. 2b), in agreement with previously-published work^28^. We also created a derivative of F1^cPPT^, designated “F1^cPPT^Δ,” in which additional coding sequence was deleted to ablate extraneous protein expression (Fig. 2a) and this construct displayed even greater increase in VLP transmission, with a three-fold increase in transduction efficiency compared to F1 (Fig. 2b).

**Figure 2:**
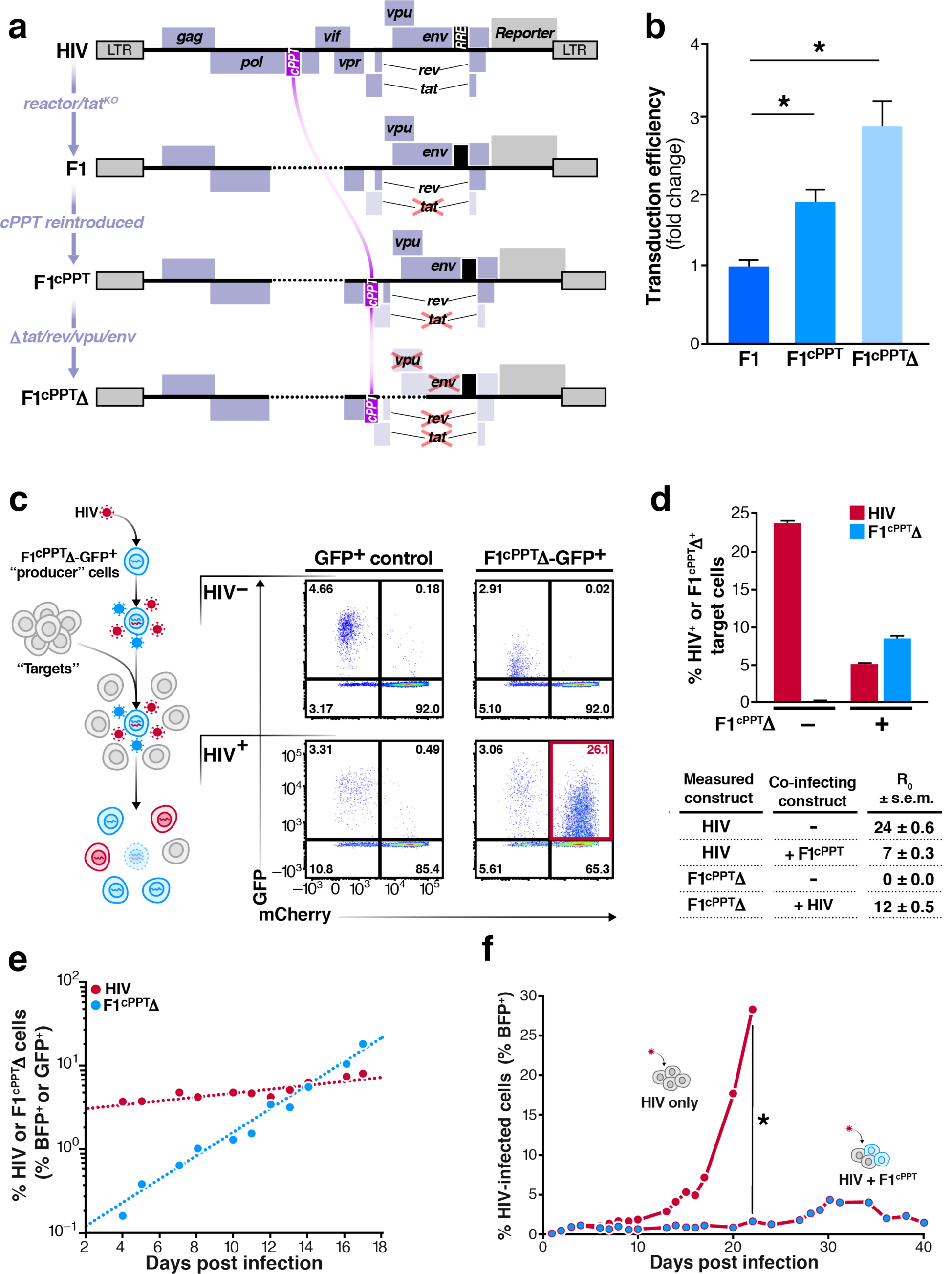
Engineering the F1 DIP into a TIP. **a**, Construct schematics for full-length HIV, the original F1 DIP construct and engineered F1 constructs. The central polypurine tract (cPPT) was added to enhance transduction efficiency in “F1^cPPT^” and protein coding sequences were deleted to reduce extraneous HIV protein expression in “F1^cPPT^Δ.” For most experiments, HIV has a BFP reporter and the F1 construct has a GFP reporter. Stop mutations were added to the first exon of Tat in F1^cPPT^ to reduce F1 transcription in the absence of co-infecting virus^54^. The first Tat exon is deleted in F1^cPPT^Δ. **b**, Transduction efficiency of optimized F1 constructs. F1 constructs, packaged as described in Fig. 1, were titered on permissive MT-4 cells and normalized to p24 (i.e., capsid protein) content. The transduction efficiency of F1 was set to 1. Data represent mean and s.e.m. of three biological replicates, *p = 0.007 and 0.003 for F1^cPPT^ and F1^cPPT^Δ, respectively, by a two-tailed, unpaired Student’s t test. **c**, Assay for mobilization of F1 constructs. Left, experiment schematic: CEM ‘producer’ cells stably transduced with F1^cPPT^Δ-GFP or a UBC-GFP control lentiviral vector were infected with HIV (or mock infected, “HIV^−^”) and at 2 dpi, co-cultured 1:20 with an MT-4 cell line transduced with an EF1A-mCherry cassette (i.e., ‘target’ cells). Right, cells were analyzed by flow cytometry 3 days post co-culture to determine whether the F1 or control constructs (GFP^+^) mobilized into mCherry^+^ target cells in the presence or absence of HIV (see red box for dual positives). **d**, R_0_ measurement of F1^cPPT^Δ. Assay setup was similar to panel c except that the target cells were CEM cells stably transduced with a dox-inducible Tat-mCherry construct. These target cells were modified to express Tat so that all transmitted F1^cPPT^Δ constructs could be detected even in the absence of co-infecting HIV. To measure R_0_, 95,000 dox-induced Tat-mCherry target cells were co-cultured with 5,000 cells that had been stably transduced with a single copy of F1^cPPT^Δ and infected with HIV-BFP. The frequencies of HIV-BFP and F1^cPPT^Δ-GFP secondary infections in these mCherry positive cells were measured by flow cytometry (top); data represent mean and s.e.m. of three biological replicates. The numbers of secondary HIV infections and F1^cPPT^Δ-transmissions, R_0_, were calculated from the frequency of BFP and GFP target cells, respectively. Mean R_0_ values ± s.e.m are shown for 3 separate experiments. **e**, Rate of HIV and F1^cPPT^Δ spread in long-term reactor culture. Spinner flasks were seeded with ∼12% F1^cPPT^Δ-GFP CEM cells and 88% mCherry^+^ target CEMs and infected with HIV-BFP at low MOI. The frequency of double-positive BFP/mCherry^+^ and GFP/mCherry^+^ cells were monitored by flow cytometry. The exponential trendlines were fit by the following equations: y(HIV) = C_1_e^0.054x^ and y(F1) = C_2_e^0.33x^. **f,** Frequency of HIV-infected cells during long-term culture in the presence and absence of F1^cPPT^-transduced cells. Cultures were seeded with a 50/50 mix of naïve cells and F1^cPPT^-transduced cells (the construct used, F1pol^cPPT^/Δenv, has an additional *env* mutation, see Supplementary Table 1) and then infected with HIV-BFP at low MOI. BFP^+^ cells were monitored by flow cytometry over time. Control culture data are omitted after 22 dpi because the culture collapsed, similar to the dynamics observed in Fig. 1c. Because trajectories represent a single culture, error bars represent technical error measurements. *p = 0.0006 by a two-tailed, paired t test.

We confirmed that both engineered constructs retained the properties of a DIP including conditional replication and packaging (i.e., they did not mobilize between cells, except in the presence of HIV), as well as interference with HIV outgrowth (Supplementary Fig. 4a-c).

To test if the new F1^cPPT^ constructs met the requirement for TIP persistence (R_0_^TIP^>1), we developed an assay to directly measure R_0_ (Fig. 2c). In brief, cells were stably transduced with a single copy of F1^cPPT^Δ-GFP (integrated at polyclonal loci) and then infected with HIV-BFP to become “producer” cells. These producer cells were then co-cultured with an excess of susceptible mCherry^+^ “target” cells so that single-round HIV and F1^cPPT^Δ transmission into target cells could be quantified from the fractions of BFP^+^/mCherry^+^ and GFP^+^/mCherry^+^ cells, respectively. Specifically, the appearance of double-positive GFP^+^/mCherry^+^ cells indicated transmission of the F1^cPPT^Δ construct from the producer cells to the target cells and the fraction of these double-positives allowed R_0_^TIP^ to be calculated (Fig. 2c; colors in the schematic do not match fluorophores to maintain consistent designation of F1 as blue and HIV as red). To test whether the assay yielded physiologically relevant R_0_ values, we measured the R_0_ of HIV alone (i.e., in the absence of F1^cPPT^Δ) and found that the value (R_0_^HIV^=24) was similar to upper estimates of R_0_ *in vivo*^22, 31, 32^. Quantifying the number of F1^cPPT^Δ transmission events after a single round of HIV infection yielded R_0_^TIP^ = 12 for F1^cPPT^Δ (Fig. 2d), satisfying the requirement for TIP persistence^3, 4, 11^.

We further verified that F1 constructs could persist over multiple rounds of infection by monitoring the transmission of the F1-GFP signal into mCherry^+^ target cells in reactor cultures. Both the F1^cPPT^Δ and F1^cPPT^ variants exhibited substantial transmission into target cells, showing an exponential rate of increase that was ∼6-fold faster than HIV over multiple rounds of infection (Fig. 2e, Supplementary Fig. 4d,e). Since F1 constructs did not affect cell growth or survival rates (Supplementary Fig. 4f-h), the increase in F1-GFP^+^ cells is likely due to efficient mobilization of the F1 constructs.

To test if the engineered F1 variant could persistently reduce viral load, we analyzed multi-week reactor infections where 50% of cells were pre-transduced with F1^cPPT^. While the infection in the control flask (100% naïve cells) increased over time, reaching a 28% infection frequency by 22 days post infection, the frequency of infection in the F1^cPPT^ culture hovered around 1-2% throughout the same timeframe (Fig. 2f). Even beyond the first three weeks, HIV infection frequency in the F1^cPPT^ culture never increased above 5%, demonstrating that F1^cPPT^ can restrict the outgrowth of HIV by about 10 fold over the course of weeks.

Next, we investigated the mechanism of F1-mediated interference. First, we tested if the F1-deletion interferes with HIV outgrowth by stealing HIV packaging resources^3, 4, 11^ (Fig. 3a, left panel). Deleting the Ψ genome packaging signal^33^ in F1^cPPT^Δ (Fig. 3b) reduced F1-associated interference by about 40% but did not completely rescue HIV titers (Fig. 3c and Supplementary Fig. 5a,b), suggesting that F1 RNA packaging provides substantial interference but that additional mechanisms must account for the remaining reduction in HIV burst size.

**Figure 3:**
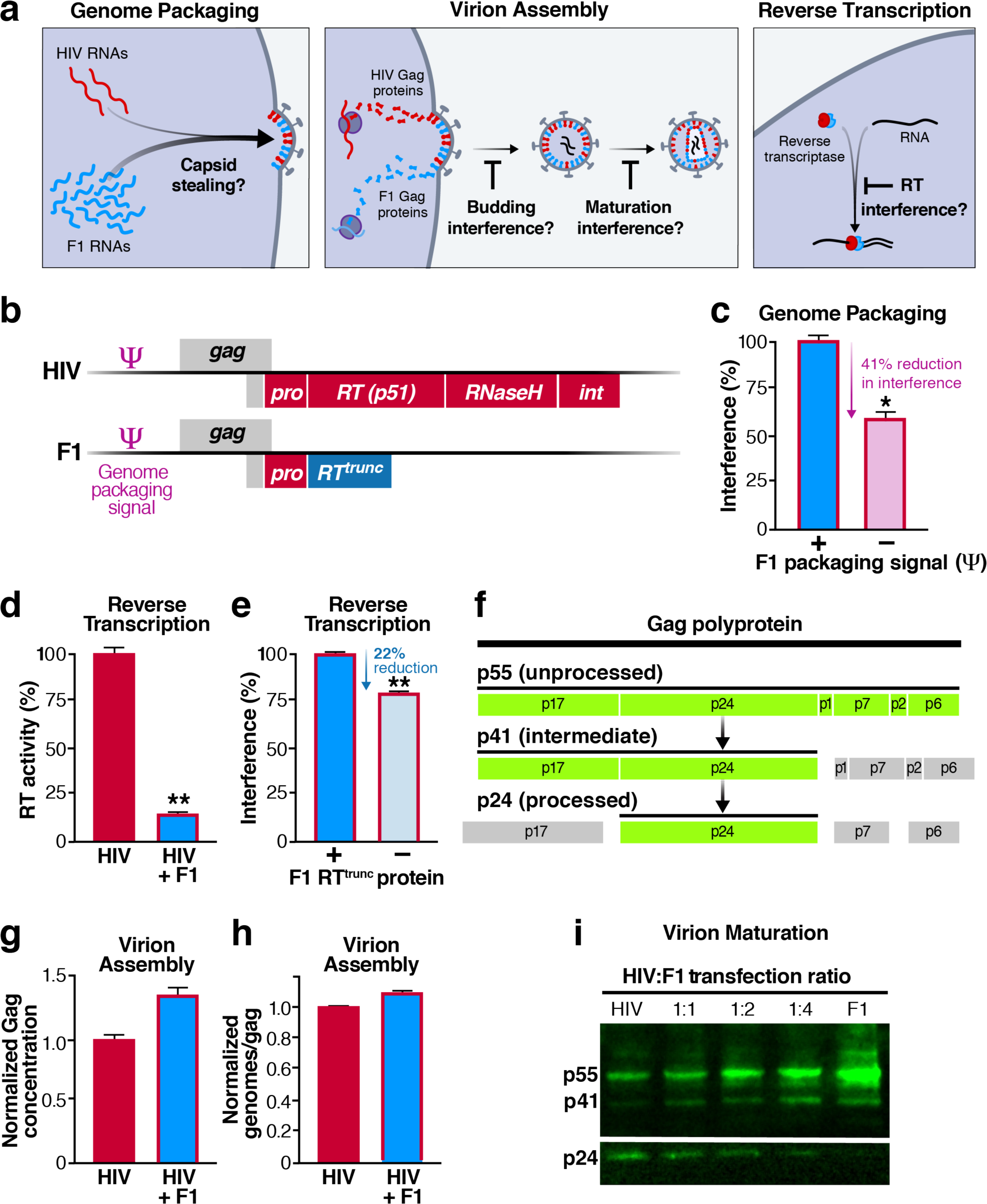
F1 interferes with HIV replication at multiple points in the viral lifecycle. **a**, Stages of the HIV lifecycle and potential targets for F1-mediated interference. Left: F1 gRNAs outcompete HIV gRNAs for access to packaging materials (i.e., “capsid stealing”). Middle: F1 proteins interfere with virion budding or maturation. Right: The F1 reverse transcriptase (RT) protein interferes with reverse transcription. **b**, Schematic of the 5’ end of the HIV and F1 constructs with the Ψ packaging signal and *gag-pol* coding regions indicated. The F1 deletion generates a truncated RT. **c**, Quantifying the interference caused by F1 genome packaging. Cells were transduced with F1^cPPT^Δ or F1^cPPT^Δ/ΔΨ (both cell populations were 50% transduced) and infected with HIV-BFP. HIV titers were measured at 3 dpi. The reduction in titer measured for F1^cPPT^Δ (52%) was set to 100% interference and the F1^cPPT^Δ/ΔΨ titer was normalized to this value. See methods for further explanation. *p = 0.009, by a two-tailed, unpaired Student’s t test. **d**, RT activity in the presence of F1. Cells transduced at high MOI with F1^cPPT^Δ (or mock-transduced controls) were infected with HIV and virions were harvested at 3 dpi. After normalizing by p24 content, RT activity of virions was measured by the SG-PERT assay^55^. **p = 0.0007, by a two-tailed, unpaired Student’s t test. **e**, Quantifying interference caused by F1 RT^trunc^ protein. An RT coding knockout mutation^34^ was introduced into the F1 construct to make F1^cPPT^Δ/ΔRT^trunc^. F1^cPPT^Δ-BFP and F1^cPPT^Δ/ΔRT^trunc^-BFP cell lines were infected with HIV-GFP. HIV titers were measured at 3 dpi and percent interference was calculated as in panel c. **p = 0.0006 by a two-tailed, unpaired Student’s t test. **f**, Schematic of the Gag proteolytic processing cascade. p24 protein is colored to indicate the species recognized by the Western blot antibody used (i.e., in panel i and Supplementary Fig. 5e and i). **g**, Total Gag protein concentration in supernatants in the presence and absence of F1. Mock or F1^cPPT^Δ-transduced cell lines were infected with HIV and Gag supernatant concentration was measured by FLAQ assay^56, 57^ at 3 dpi. **h**, Ratio of packaged genomes to Gag protein in the presence and absence of F1. Total virion RNA (HIV + F1^cPPT^Δ) was quantified by RT-qPCR and normalized to Gag protein concentration measured in panel g. **i**, Western blot analysis of Gag proteolytic processing at various ratios of HIV and F1. Plasmids encoding full-length HIV or F1(Tat+) (to drive protein expression) were co-transfected into 293T cells at the indicated molar ratios. Supernatants were harvested, purified by ultracentrifugation and Western analysis was performed with anti-p24 antibody (to compare higher molecular weight bands to lower molecular weight bands in the same panel the image was cropped: top of image is from a shorter exposure while bottom of image (i.e., p24 bands) is from a longer exposure).

Since the F1 deletion truncates the reverse transcriptase (RT) gene (“RT^trunc^,” Fig. 3b) and pure F1 VLPs do not display RT activity (Supplementary Fig. 5c), we tested if the presence of F1 reduced RT activity in virions released from HIV-infected cells. Not only did F1^cPPT^Δ reduce virion-associated RT activity (Fig. 3d), RT^trunc^ proteins were found in purified F1 VLPs (Supplementary Fig. 5d,e) suggesting that defective RT^trunc^ may compete with wild-type RT for virion access and/or RNA templates and subsequently fail to complete reverse transcription (Fig. 3a, right panel). Knocking out translation^34^ of RT^trunc^ in either the F1^cPPT^ or F1^cPPT^Δ background reduced F1-associated interference by 22% (Fig. 3e, Supplementary Fig, 5f), suggesting that the RT^trunc^ protein also contributes to the reduction in HIV titers.

Having accounted for a portion of F1-mediated interference, the literature suggested a third possible avenue for interference: protease defects. Truncations in the RT sequence are known to reduce protease activity^35^ and catalytically-dead proteases *trans-*dominantly interfere with virion maturation by failing to cleave the Gag structural proteins^36^ (Fig. 3f, Gag cleavage cascade). While the presence of F1 did not affect HIV virion assembly and budding—supernatant concentrations of Gag (Fig. 3g and Supplementary Fig. 5g) and the gRNA:Gag ratio (Fig. 3h and Supplementary Fig. 5h) were unchanged—Western blot analysis revealed a severe defect in Gag proteolytic cleavage (Fig. 3i), similar to that observed in catalytically dead protease mutants (Supplementary Fig. 5i). Consistent with the fact that Gag processing defects are potent dominant negatives^37, 38^, the presence of wild-type protease did not fully rescue the Gag processing defect (see lanes 2-4 in Western blot, Fig. 3i), suggesting that F1 may interfere with HIV transmission by arresting virions in the immature, non-infectious state (Fig. 3a, middle panel).

In sum, F1 appears to interfere with the outgrowth of HIV via at least three distinct points in the viral lifecycle: genome packaging, virion maturation, and reverse transcription, mimicking a combination therapy. Some of the interference appears to involve dominant-negative interactions, which are reported to have a high barrier to therapeutic resistance^39, 40^. Given that F1^cPPT^ and F1^cPPT^Δ exhibit similar packaging efficiency, HIV interference levels, (Supplementary Fig. 4a-c), and interference with RT (Fig. 3e vs Supplementary Fig. 5f), the relative breakdown of F1-mediated interference mechanisms is likely not specific to a particular F1 construct.

We next tested the therapeutic effect of the engineered F1 constructs in a humanized mouse model^41, 42^ used in preclinical trials for new HIV therapies. In this model, hu-PBL mice are engrafted with human CD4+ T cells and subsequently infected with HIV. CD4+ T cell counts decline over the course of a few weeks due to virally-induced cell death, and we hypothesized that mice engrafted with F1-transduced cells should maintain their CD4+ T cell population compared to controls (see Fig. 4a for experimental timeline). As expected, CD4+ T cell counts declined in HIV-infected mice within two weeks. However, mice that received F1^cPPT^ or F1^cPPT^Δ-transduced cells retained their CD4+ T cell population (Fig. 4b, p ≤ 0.003, see Supplementary Fig. 6a for gating scheme). The T cell protection in F1-treated mice correlated with a one-log suppression in viral load that occurred a week prior to the observed difference in T cell counts (Fig. 4c, p ≤ 0.0001), suggesting that the lower viral load prevented T cell depletion. F1-associated T cell maintenance was also observed in an independent cohort, confirming the therapeutic effect of the F1 construct (Supplementary Fig. 6b). We also observed F1 RNAs in the serum (Fig. 4d), indicating that engineered F1 constructs mobilized within the mouse. In some animals, F1 RNAs accounted for greater than 50% of serum RNAs despite the fact that only ∼50% of engrafted cells were transduced with F1 constructs. These data suggest that F1 constructs may be a viable avenue to suppress HIV, though long-term *in vivo* efficacy will require further testing.

**Figure 4:**
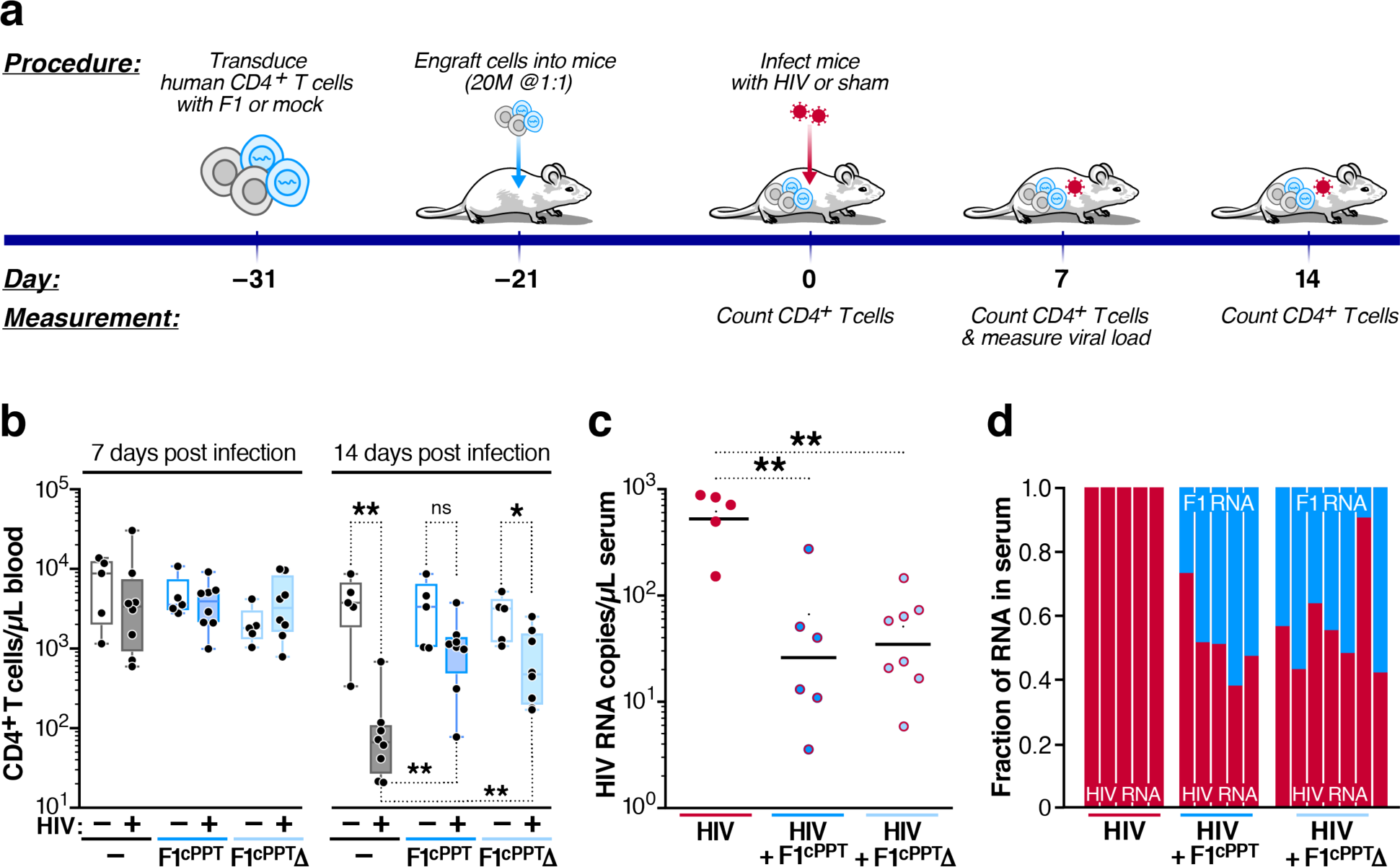
F1 lowers HIV viral loads and protects CD4+ T cells in a humanized mouse model. **a**, Experimental timeline for engraftment, infection and analysis of hu-PBL mice engrafted with primary human CD4+ T cells. **b**, CD4+ T cell concentration in F1-treated and control mice, with and without HIV infection at one (left) and two (right) weeks post infection. The F1^cPPT^Δ construct in this experiment is the F1stop^cPPT^Δ construct which contains a double stop codon early in the F1 RT coding region but was not found to alter F1 interference in the mouse (see Supplementary Fig. 6b for F1^cPPT^Δ data). Box and whiskers plots, n = 8 for each condition. Black dots represent individual mice, line represents the median, the box extends from the 25^th^ to 75^th^ percentile and whiskers denote the maximum to minimum values. Asterisks at top of graph indicate p values calculated within group (i.e., +/- infection) while asterisks at bottom of the graph indicate p values calculated between infected groups (e.g., control [no F1^cPPT^] vs F1^cPPT^ mice). P values were calculated using the Mann-Whitney test and significant values are as follows: control mice +/- infection, 0.003; F1^cPPT^Δ mice +/- infection, 0.03; infected control vs infected F1^cPPT^ mice, 0.002; infected control vs infected F1^cPPT^Δ mice, 0.003. *NB:* one mouse from each infected F1 group was euthanized due to graft versus host disease – a symptom of high levels of CD4+ T cells – two weeks post infection. **c**, Viral loads in mouse serum collected from infected mice one week post infection were measured by digital droplet PCR (ddPCR). Points represent individual mouse samples and line represents the geometric mean. P values were calculated using a one-way ANOVA followed by Tukey’s multiple comparisons test and were as follows: control vs F1^cPPT^ mice, p = 0.0001; control vs F1^cPPT^Δ mice, p < 0.0001. Three samples from control mice and two samples from F1^cPPT^ mice were lost during RNA purification. **d**, Fraction of full-length HIV and F1 gRNA in mouse serum one week post infection. Each bar represents a single HIV-infected mouse. Mice with viral loads lower than 10 copies/μl were excluded from analysis.

Overall, the data presented here provide proof-of-concept for a class of biologic that can conditionally self-renew and thereby has the potential to be developed into a single-administration therapeutic. The data indicate that F1^cPPT^ variants meet the definition of a TIP: they self-replenish by conditionally transmitting between cells with R_0_>1, display sustained inhibition of HIV in culture, and prevent CD4+ T cell depletion in a pre-clinical humanized-mouse model. Despite only ∼1% of cells being productively HIV infected at any given time during asymptomatic infection *in vivo*^22, 43^, established HIV dynamics models indicate that the rapid daily turnover of this 1% of HIV-infected cells would enable TIPs with an R_0_>1 to co-infect sufficient numbers of cells thereby reducing viral set point by 1–2 Logs^3, 4, 10, 11^—similar to rationales explaining rapid HIV recombination despite most cells harboring a single HIV provirus^44^.

A potential criticism is that TIPs, unlike antiretroviral therapies, do not reduce HIV’s *in vivo* R_0_ below 1. However, models indicate that if TIPs forced HIV’s R_0_ below 1, TIPs would be unable to maintain R_0_>1 and therefore would not persist, allowing HIV to reestablish pre-treatment viremia levels after rebound from latency^3, 4, 10, 11^. The models further predict that TIPs with R_0_>1 would possess unique therapeutic capacities including the ability to efficiently access difficult-to-reach high-risk populations, surmount behavioral and adherence barriers, and circumvent mutational escape by establishing co-evolutionary arms races with HIV. Notably, epidemiological models^4, 10^ indicate that despite antiretroviral therapy blocking TIP activity at the individual-patient scale, at the population scale TIPs would primarily target high-risk groups where therapy coverage is low and would thus compliment ongoing antiviral campaigns.

Clearly, the long-term efficacy of TIPs will need to be addressed, including whether the R_0_ is sufficient to control viral loads in a reasonable amount of time *in vivo*, when and how TIPs should be introduced *in vivo* for maximal efficacy, and the co-evolutionary dynamics of HIV and TIPs^45^. In addition, the safety of TIPs—both genotoxicity due to TIP integrations and immunotoxicity due to TIP viral loads—will need to be experimentally addressed. We and others have speculated that TIPs may be no more detrimental in patients than the large reservoir of defective proviruses^46, 47^ since TIPs are constructed solely from sequences already found in HIV and, regarding genotoxicity, clinical studies indicate that integrating lentiviral gene therapies do not generate an increase in T cell cancers^48–50^. While recombination between HIV and TIP (or existing defective proviruses) is a potential concern and will need to be thoroughly examined, the large internal deletion in F1 reduces the efficiency of recombination, and the most likely recombination product is replication-competent HIV^11^, which will only be generated if HIV is present and would thus be equivalent to a conventional loss of therapeutic efficacy. Future studies will determine if the ability of F1^cPPT^ vectors to simultaneously interfere with HIV and enhance their own VLP production are separable properties, and if the potential for independent genetic re-assortment of these properties exists.

From the standpoint of evolutionary dynamics, one question is why F1-like TIPs have not been detected *in vivo* despite the well-known abundance of defective HIV genomes^46, 47^ or previously selected *in vitro*. A potential explanation could be that for such deletions to accumulate requires an R_0_>1, which necessitates an unlikely series of recombination and genome-rearrangement events where the cPPT is retained or reinserted whilst specific flanking sequences are deleted^47^.

Finally, the reactor selection result (Fig. 1c), where host cells ultimately recovered from HIV infection by integrating a defective viral sequence into their genomes (a phenomenon we have observed repeatedly in culture), is particularly intriguing. Similar *in vitro* reactor systems may provide a model to recapitulate and examine real-time host cooption or “exaptation” of endogenized retroviral sequences for antiviral defense^51^, or protein repurposing that is common to host-pathogen arms races^52, 53^. Ultimately, these selection schema may enable discovery of defective variants that can be reengineered into countermeasures that suppress viral resistance, echoing recent advances in phage therapy^2^.

---

**Supplementary Figure 1:**
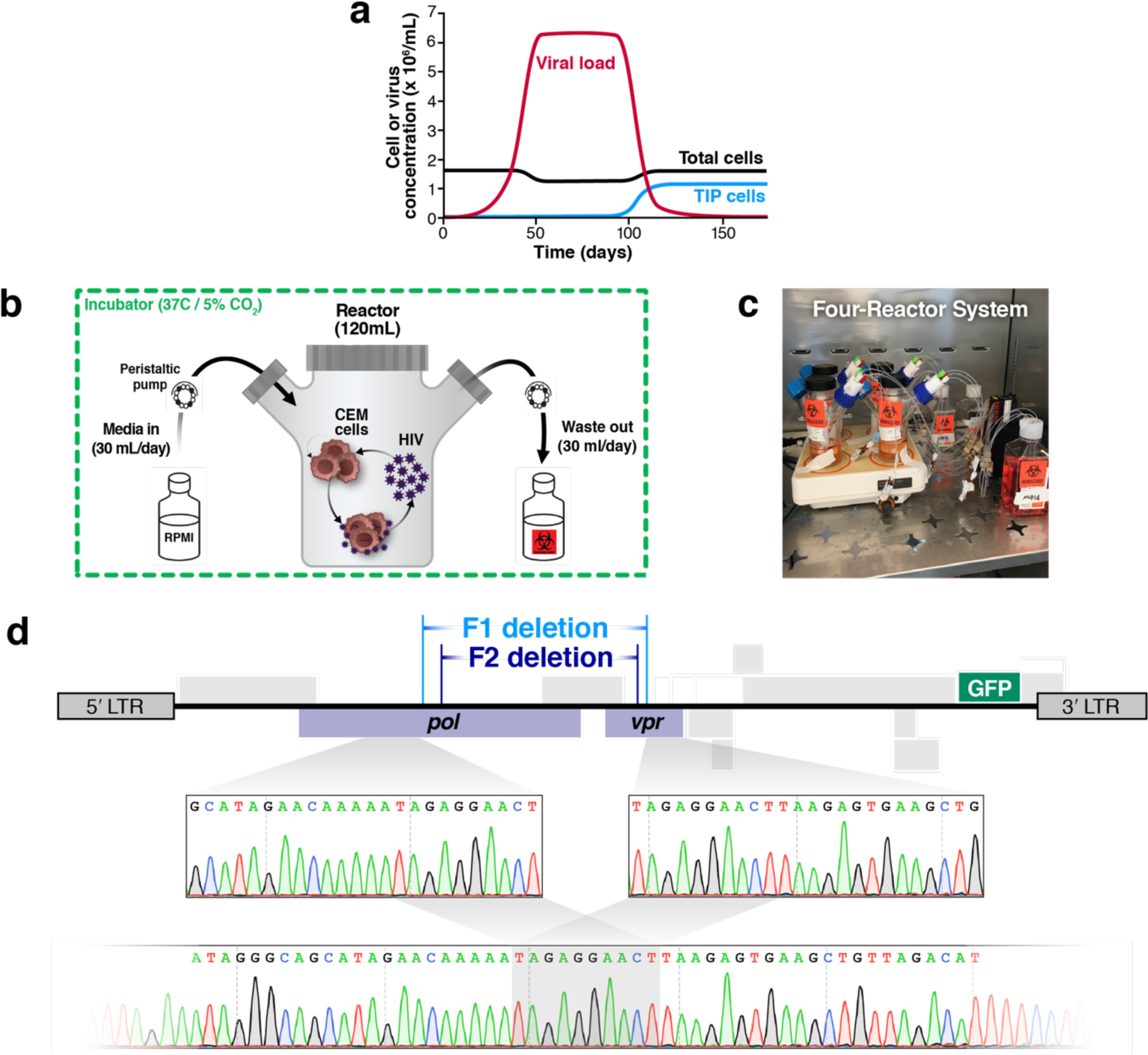
Selection for HIV DIPs in long-term culture; mathematical model prediction, reactor schematic, and deletion variant sequencing. a, Numerical simulations (see Supplementary Methods) of a computational model of HIV dynamics in a long-term culture where cells are replenished through cell division. Modeling predicts that if a DIP spontaneously arises on day one of the culture, it will take ∼100 days to expand to appreciable levels. b, Cartoon of closed-system reactor setup. c, Photo of a reactor set up where four reactor flasks are being run in parallel. d, Deletion variants detected in two parallel long-term reactors. The location of each deletion is indicated above a schematic of the original HIV-GFP genome that was used to infect the long-term culture. The F1 deletion spans nucleotides 3159-5630 and the F2 deletion spans nucleotides 3973-5556 (relative to the NL4-3 proviral sequence coordinates). Below the genome schematic, the junction of the F1 deletion is indicated, including the sequence of 10 base-pair homology regions.

**Supplementary Figure 2:**
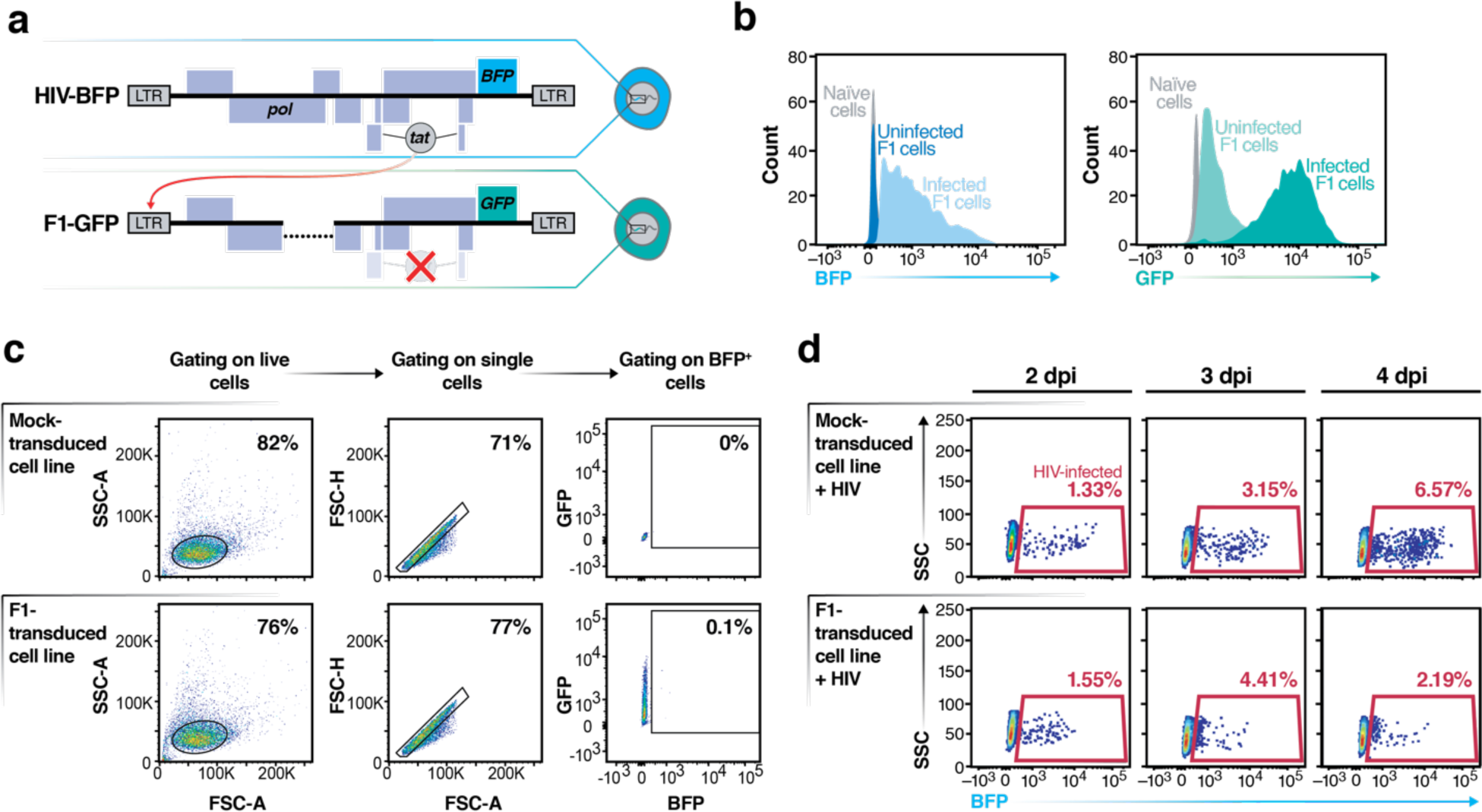
The F1 construct is transcribed during HIV infections and inhibits HIV spread. a, Schematic of HIV-BFP (the NL4-3 molecular clone in which the *nef* reading frame is replaced with *BFP-IRES-nef*) and the F1-GFP construct. In F1, two stop codons were introduced into the first exon of Tat^54^, the HIV transactivator protein that drives high level transcription from the LTR promoter. High levels of F1 transcription should only be achieved when Tat is provided *in trans* by HIV coinfection. The F1 construct was packaged similarly to a lentiviral vector and integrated into cells via transduction (see methods). b, HIV infection of an F1-GFP cell line activates GFP expression. Left: BFP (i.e., HIV) expression +/- HIV infection; Right: GFP (i.e., F1) expression +/- HIV infection. The F1-GFP cell line was created by transducing the F1 construct in panel a into CEM T cells at low MOI (less than 15% transduced) and batch sorting cells with dim GFP expression. c, Flow cytometry dot plots showing representative gating scheme on HIV-negative samples. d, Flow cytometry dot plots of HIV-infected BFP^+^ cells at 2, 3, and 4 days post infection. Mock transduced (control) or F1 cells (polyclonal, single integrations) were infected with HIV-BFP at low MOI and analyzed by flow cytometry.

**Supplementary Figure 3:**
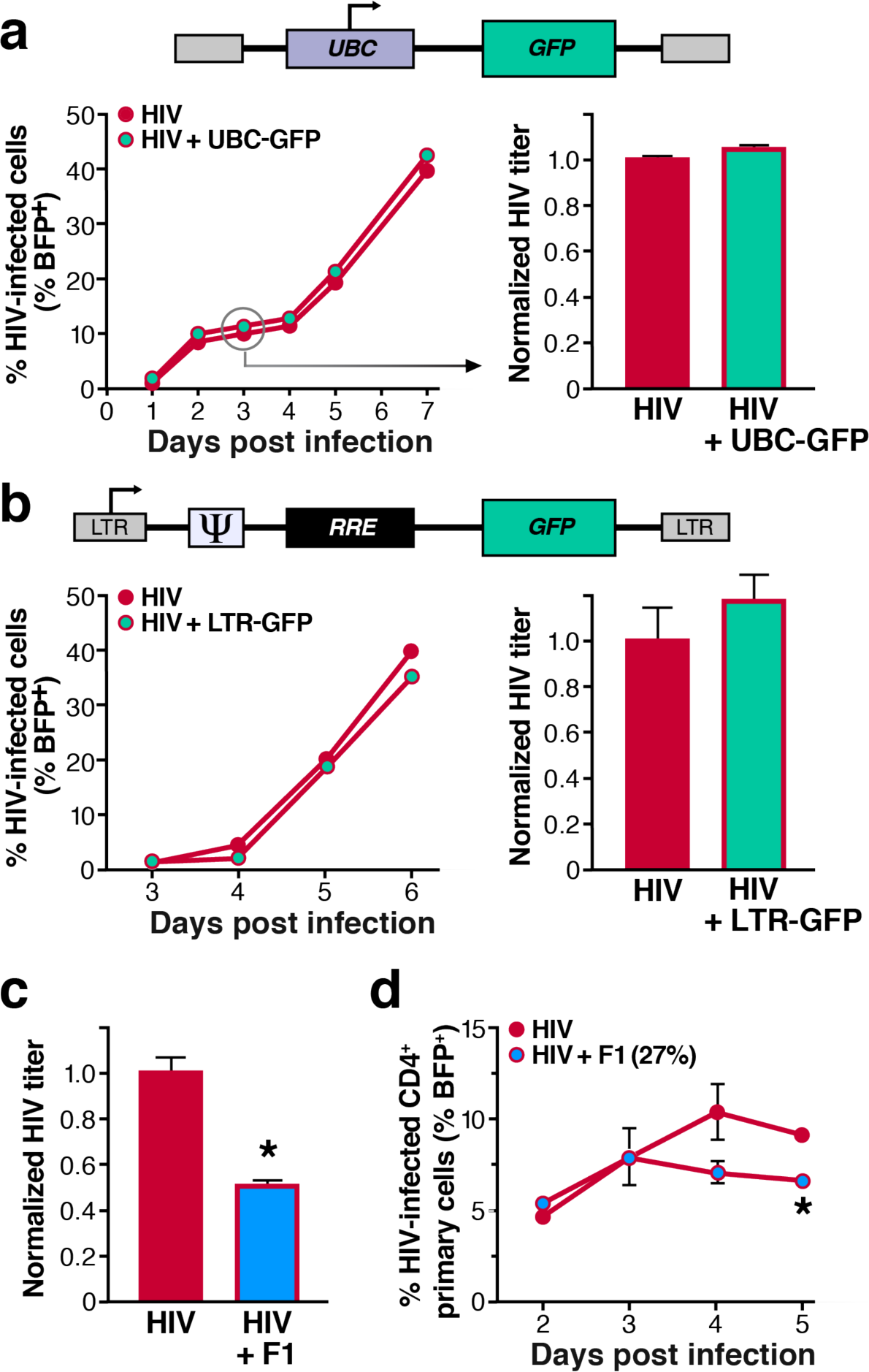
The F1 construct interferes with HIV outgrowth in various cell types while other lentiviral vectors do not. a, Multiple and single-round HIV outgrowth in the presence of a control lentiviral vector. Top, schematic of the integrated lentiviral vector which expresses GFP from the UBC promoter (pFUGW^58^). Empty gray boxes represent lentiviral LTRs that enable vector integration. Cell lines were created as in Supplementary Fig. 2b except that GFP expression from the UBC promoter was bright rather than dim. Left panel, the frequency of HIV-infected cells (i.e., BFP^+^) was determined by flow cytometry at various days post infection. Right panel, single round titers were determined by infecting cells with HIV-BFP at high MOI (81% infected at 3 dpi), harvesting filtered supernatant at 3 dpi and titering on a MT-4s by flow cytometry. b, Multiple and single-round HIV outgrowth in the presence of a lentiviral vector that harbors sequence elements involved in HIV genome packaging (Ψ) and nuclear export (RRE). Similar to F1, the LTR acts as the promoter in this construct (pLG^24^). Multi- and single-round HIV infections were analyzed as in panel a except that CEMs were transduced with the construct but not sorted. Left panel, 67% of cells were transduced with LTR-GFP while data in right panel were obtained from cells that were 81% transduced with LTR-GFP and 26% infected with HIV-BFP. c, Single-round HIV titers in the presence of F1 in MT-4 cells. An F1-GFP MT-4 cell line was created as described in Supplementary Fig. 2b and infected with HIV-BFP at high MOI (81% of cells were infected at 3 dpi). 3 dpi supernatant titers were determined by flow cytometry. *p = 0.002 by a two-tailed, unpaired Student’s t test. d, Spread of HIV in human primary CD4+ T cells in the presence and absence of F1. Human primary CD4+ T cells were co-spinoculated with HIV-BFP and F1-GFP (or mock) and % BFP^+^ cells was measured by flow cytometry at various days post infection. Only 27% of cells were co-infected/transduced with HIV and F1 at 3 dpi. On day 4, p = 0.08 and on day 5, *p = 0.002 by a two-tailed, unpaired Student’s t test. Data represent the average and s.e.m. of three biological replicates.

**Supplementary Figure 4:**
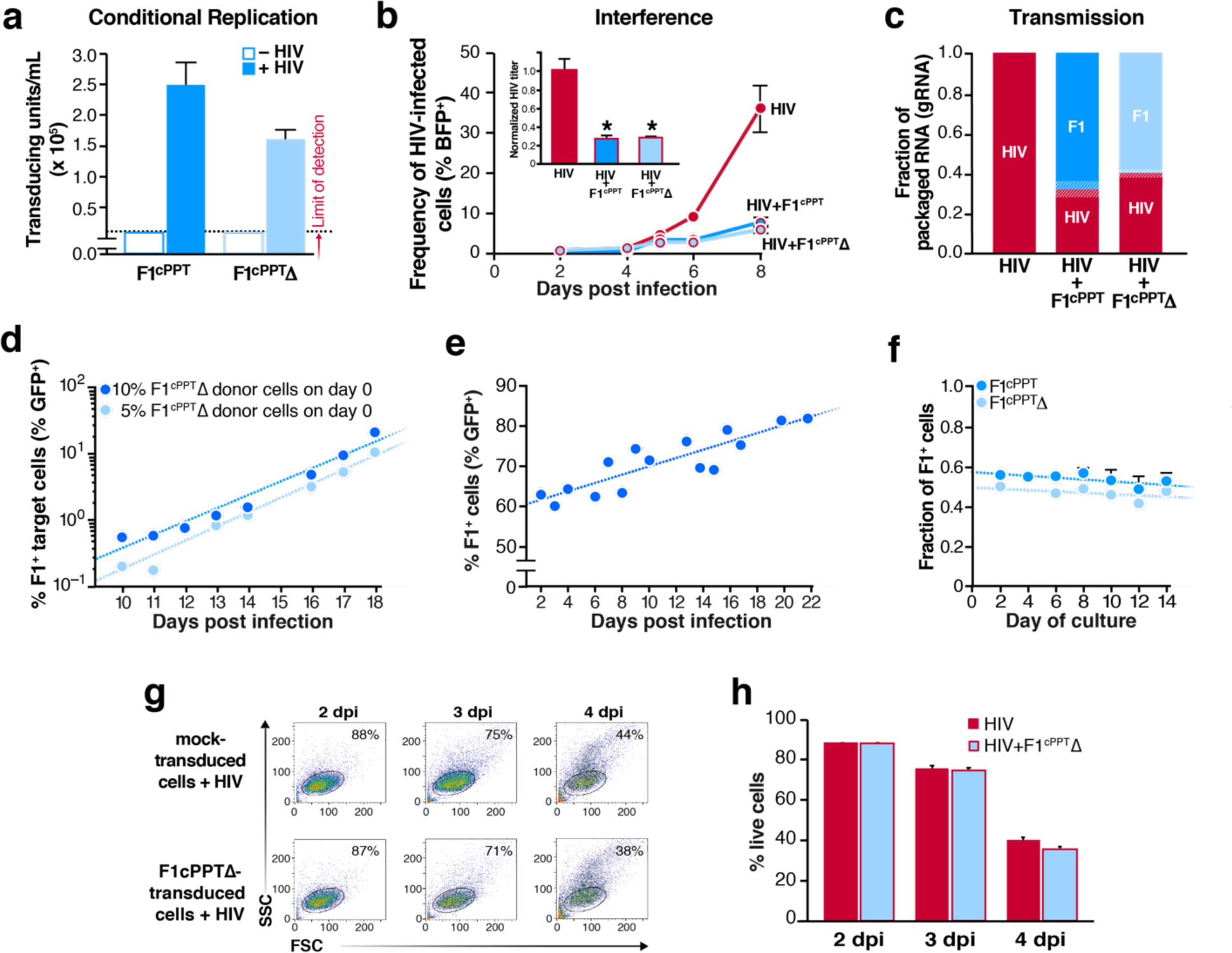
Engineering F1 to increase transmission for long-term control of HIV outgrowth. a, Transduction competence of engineered F1 VLPs assembled in the presence and absence HIV. Polyclonal cell lines transduced with engineered F1 constructs (single integrations with GFP reporter) were infected with HIV and supernatant was harvested at 3 dpi. F1^cPPT^ and F1^cPPT^Δ VLPs were tittered on permissive MT-4 cells by flow cytometry for GFP. b, Outgrowth of HIV in the presence and absence of the engineered F1 constructs. Polyclonal F1^cPPT^-GFP or F1^cPPT^Δ-GFP cell lines (same as in a) or a mock-transduced control were infected with HIV-BFP at low MOI. The frequency of BFP^+^ cells was measured by flow cytometry at indicated timepoints. P values were calculated by a two-tailed, unpaired Students t test. HIV only vs +F1^cPPT^: p = 0.002 (6 dpi), p = 0.004 (8 dpi); HIV only vs +F1^cPPT^Δ: p = 0.0009 (6 dpi), p = 0.02 (8 dpi); Inset: Filtered supernatants, harvested at 3 dpi, were titered by flow cytometry (48 after infection of naïve cells). p = 0.005 for titers of HIV only vs each F1 construct by a two-tailed, unpaired Students t test. c, The fraction of packaged RNAs that are derived from HIV or engineered F1 variants. Samples titered in panel b (inset) were analyzed by allele-specific RT-qPCR. d, The frequency of transmitted F1^cPPT^Δ in cultures established with different starting frequencies of F1^cPPT^Δ cells. Spinner flasks were seeded with ∼10% or ∼5% F1^cPPT^Δ CEMs, ∼14% mCherry^+^ target CEMs and untrasduced CEMs made up the remainder of the cell population. F1^cPPT^Δ-GFP mobilization was indicated by the percent of GFP^+^ cells in the mCherry^+^ gate. Exponential trendlines were added using Microsoft Excel. e, The frequency of F1^cPPT^-GFP cells in a reactor infection. The frequency of total GFP^+^ cells was monitored by flow cytometry in the long-term culture experiment shown in Fig. 2f. mCherry^+^ target cells were not included in this reactor so the total frequency of GFP^+^ cells (sum of seeded F1^cPPT^ cells + transmission events) is reported. Linear trendline was added using Microsoft Excel. f, The fraction of engineered F1 cells in co-culture with mock-transduced CEMs over time. F1^cPPT^ or F1^cPPT^Δ cell lines were mixed 1:1 with mock-transduced cells and their frequency was measured over time. Linear trendline was added using Microsoft Excel. g, Representative flow cytometry scatter plots of mock-transduced or F1^cPPT^-transduced CEM cells following HIV infection at an MOI of ∼1. The frequency of cells falling in the live gate is noted on the plot. h, Quantification of cells in the live gate including data shown in panel g. Data in panels a,b,c,f and h represent mean and s.e.m of three biological replicates. Data in panels d and e are from individual culture trajectories and so do not have biological repeats or error bars.

**Supplementary Figure 5:**
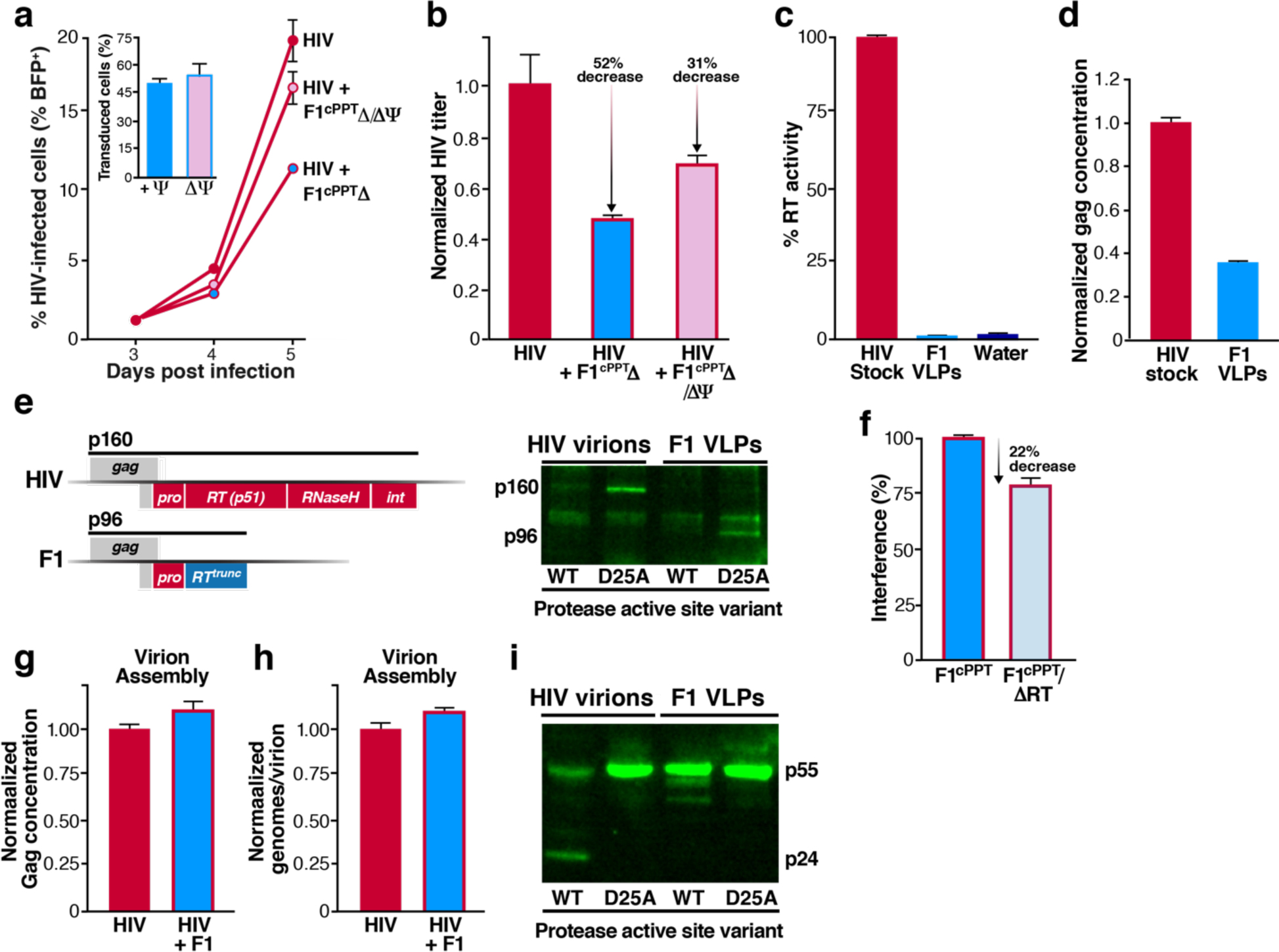
F1 constructs interfere with HIV replication combinatorially through capsid stealing, defective reverse transcriptase activity, and a gag processing defect. a, Outgrowth of HIV in the presence of packaging-incompetent F1. CEM cells were transduced with F1^cPPT^Δ or F1^cPPT^Δ/ΔΨ so that ∼50% of cells harbored the F1 constructs (see inset). Cells were infected with HIV-BFP and the frequency of BFP^+^ cells was measured by flow cytometry at indicated timepoints. b, Relative HIV titers in the absence and presence of packaging-incompetent F1 (related to Fig. 3c). Assay setup as in panel a except that supernatants were harvested at 3 dpi and titered on the permissive MT-4 cell line. c, Reverse transcriptase activity of pure F1 VLPs. Virions were harvested from 293T transfections with plasmids encoding full-length HIV or F1(Tat+) (the Tat coding sequence was left intact to drive protein expression). HIV and F1 stocks were normalized for capsid protein content and RT activity was measured by the SG-PERT assay^55^. SG-PERT uses virion-associated RT proteins to reverse transcribe a template RNA and the cDNA is quantified by qPCR. Here, the Ct values were converted to fold change relative to HIV stock, which was set at 100% RT activity. d, Gag concentration for stocks of pure HIV virions or F1 VLPs. Samples are same as those in panel c prior to normalization for Gag protein content. e, Western blot analysis of F1 GagPol protein packaging. Left: Schematic of wild-type and F1 *gag-pol* coding regions with total polyprotein size indicated. Right: Western blot of GagPol proteins in the presence of wild-type protease or a catalytically inactive protease mutant (D25A). 293T cells were transfected with full-length HIV or F1(Tat+) plasmids with and without catalytically active protease domains and supernatant virions were purified by ultracentrifugation before Western analysis with an antibody to p24. The catalytically inactive protease mutation (D25A)^59^ enables detection of the GagPol precursors in virions using an antibody to p24. f, Quantifying interference caused by F1 RT^trunc^ protein (related to Fig. 3e). An RT coding knockout mutation^34^ was introduced into the F1^cPPT^ construct to make F1^cPPT^/ΔRT^trunc^. F1^cPPT^-BFP and F1^cPPT^/ΔRT^trunc^-BFP cell lines were infected with HIV-GFP. HIV titers were measured at 3 dpi and percent interference was calculated as in Fig. 3c *p = 0.003 by a two-tailed, unpaired Student’s t test. g, Gag protein concentration in supernatants in the presence and absence of F1. Mock-transduced or F1 cell lines were infected with HIV and the Gag supernatant concentration was measured by FLAQ assay^56, 57^ at 3 dpi. The value obtained for infection of mock-transduced cells was set to 1. h, Ratio of packaged genomes to Gag protein in the presence and absence of F1. Total virion RNA (HIV + F1) was quantified by RT-qPCR and normalized to the Gag protein concentration measured in panel g. The values obtained for infection of mock-transduced cells was set to 1. i, Western blot analysis of Gag proteolytic processing defect. Purified virions produced with and without catalytically active protease were analyzed with an antibody to p24.

**Supplementary Figure 6:**
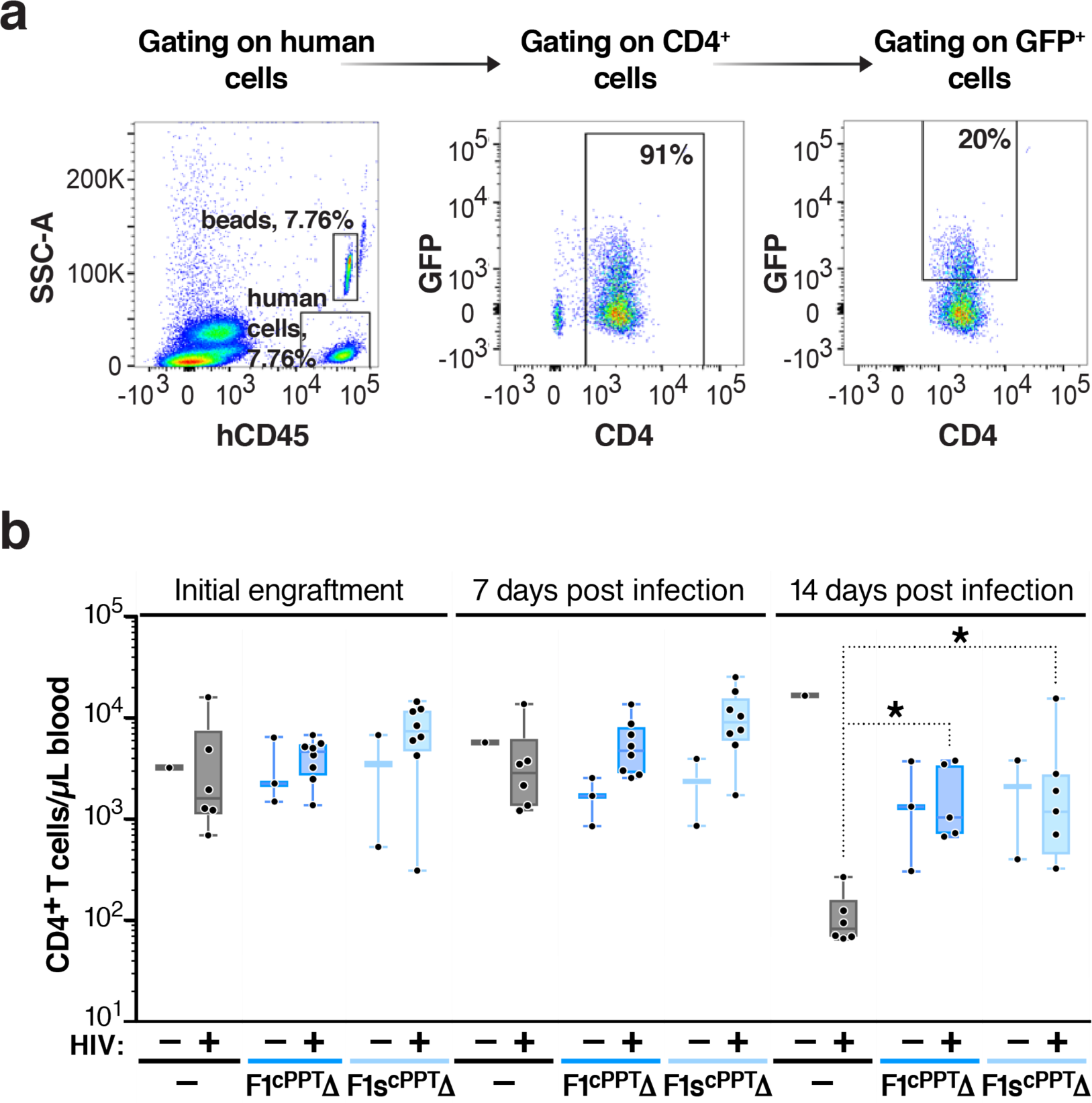
F1 constructs lower HIV viral loads and protect CD4+ T cells in a humanized mouse model. a, Gating scheme for human CD4+ T cell counts. b, CD4+ T cell concentration in F1-treated and control mice, with and without HIV infection. Data are shown prior to infection (left) and at one (middle) and two (right) weeks post infection. Numbers of mice in each group were as follows: mock transduced: 1 HIV-, 6 HIV+; F1^cPPT^Δ: 3 HIV-, 8 HIV+; F1stop^cPPT^Δ: 2 HIV-, 8 HIV+. F1stop^cPPT^Δ contains a double stop codon early in the F1 RT coding region (see constructs table). Box and whiskers plots; black dots represent individual mice, line represents the median, the box extends from the 25^th^ to 75^th^ percentile and whiskers denote the maximum to minimum values. P values were calculated using the Mann-Whitney test and significant values are as follows: infected control vs infected F1^cPPT^Δ mice, 0.004; infected control vs infected F1stop^cPPT^Δ mice. *NB*: two mice from each infected F1 group were euthanized due to graft versus host disease (GVHD)—a symptom of high levels of CD4+ T cells due to protection from HIV infection—at week one or two post infection.

**Supplementary Table 1:**
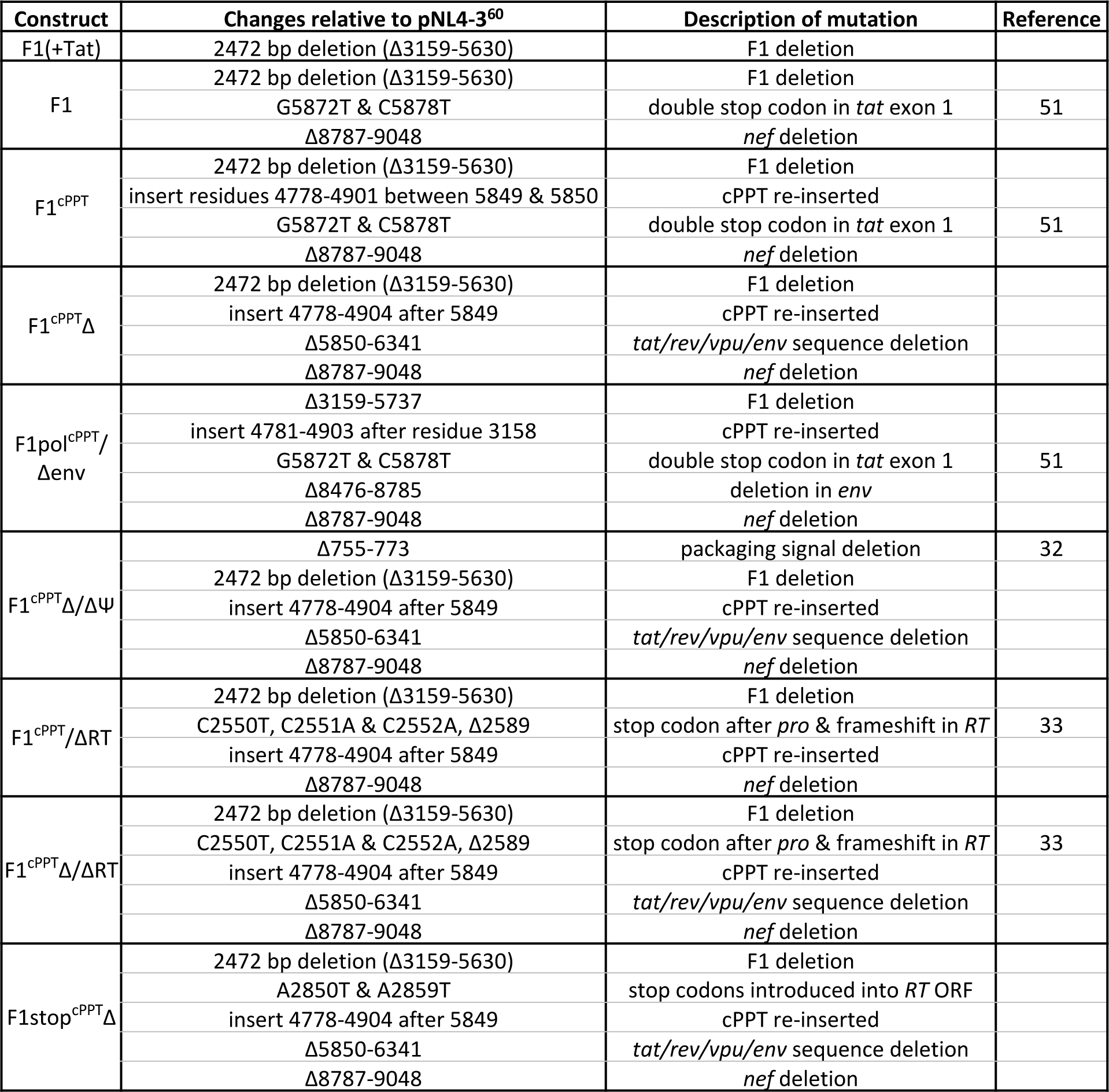
Constructs created for this work

**Supplementary Table 2.**
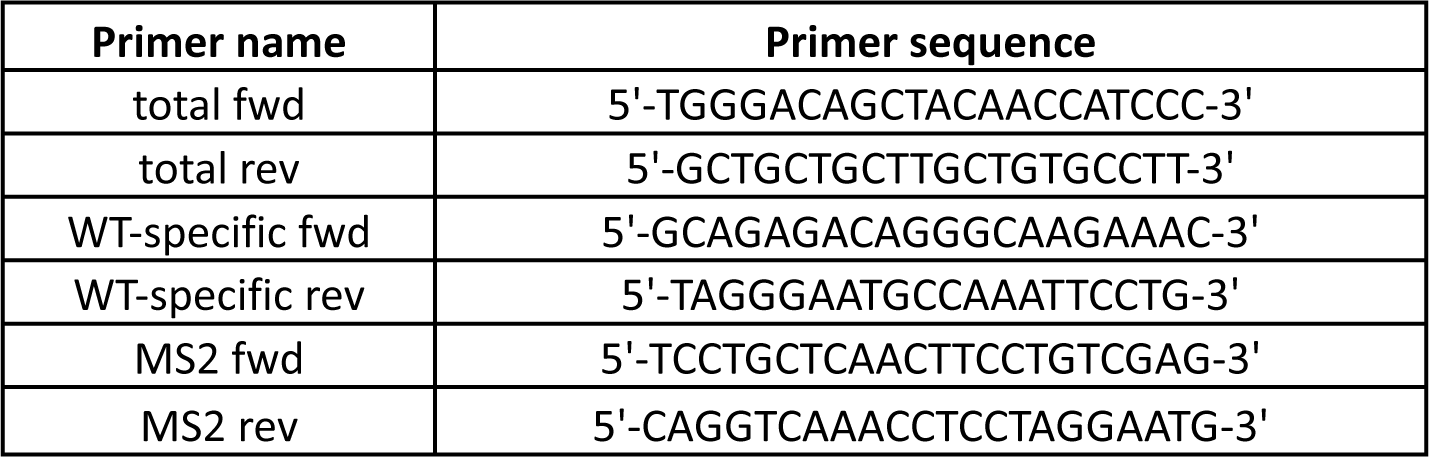
: qPCR primers

**Supplementary Table 3.**
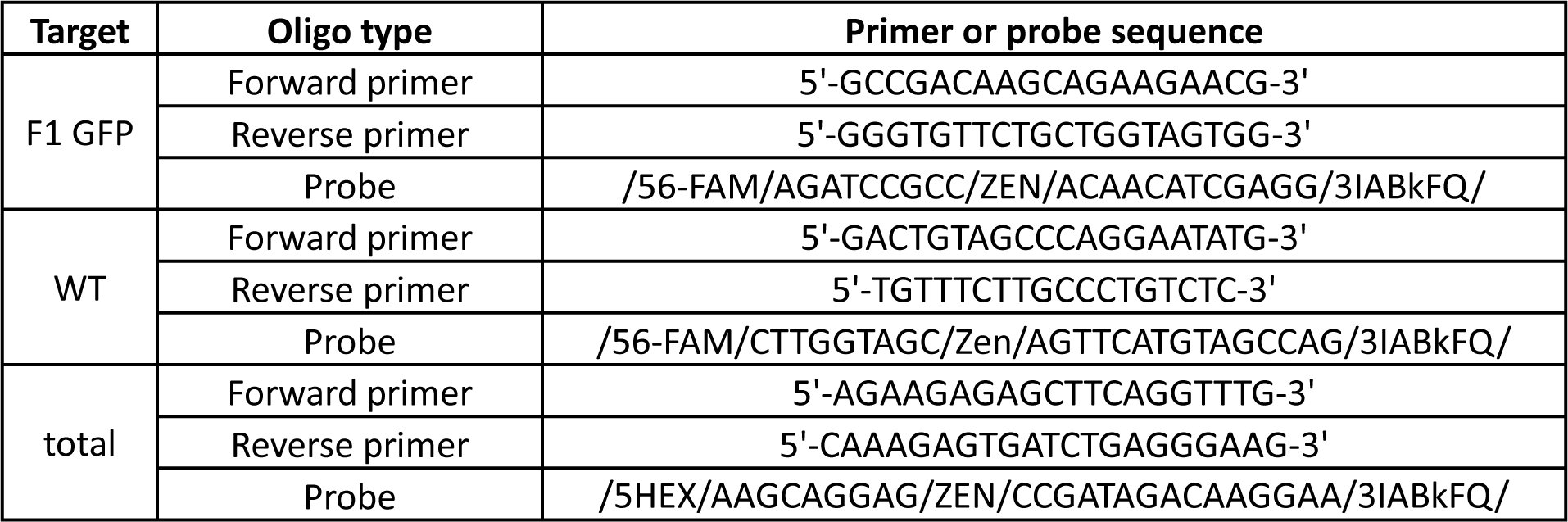
: digital droplet PCR primers and probes

## Materials and Methods

### Cell lines

All cells were grown at 37°C with 5% CO_2_. MT-4 (NIH AIDS Reagent Program, 120) and CEM CD4+ (NIH AIDS Reagent Program, 117)^60^ cells were maintained in RPMI 1640 (Fisher Scientific, MT10040CV) supplemented with 10% FBS (Fisher Scientific, MT35011CV) and 1% Pen/Strep (Fisher Scientific, MT30002CI). 293Ts (ATCC, CRL-3216) were maintained in DMEM (Fisher Scientific, MT10013CV) supplemented as for RPMI.

### Long-term culture infection and analysis

Once reactors were constructed (see supplementary information), cells were added at ∼0.5 – 1 million cells/mL in 80-120 mL volumes and cultures were monitored for cell density and viability for one to two weeks to ensure stable cell concentration and viability. Once cultures were stable, cells were infected with replication competent HIV (with either a GFP or BFP reporter) at low MOI (∼0.01 – 0.1, depending on experiment). On indicated days post infection, 3 mL of total flask contents (i.e., cells and supernatant) were removed from flasks and analyzed. Cell concentration and viability were determined using a Bio-Rad TC20 Automated cell counter (Catalog #1450102), cells were fixed in 1% formaldehyde (Tousimis Research Corporation, 1008A) and fluorescence was measured on a BD LSR II flow cytometer and analyzed in FlowJo.

### PCR and Sanger sequencing of proviral DNA

Total genomic DNA was extracted from about 1 million long-term culture cells using the NucleoSpin Tissue kit (740052.50) according to the manufacturer’s protocol. PCR was performed with the following previously published primers, modified for NL4-3 sequence^61^: 5’-CCTCAGATCACTCTTTGGCAGCGAC-3’ and 5’-CTCTCATTGCCACTGTCTTCTGCTC-3’. DNA was amplified using Phusion High-Fidelity PCR Master Mix (NEB, M0531S) and the following conditions: initial denaturation: 98° C for 30 seconds; cycle 30 times: 98° C for 10 seconds, 60° C for 30 seconds, 72° C for 4 minutes; final extension: 72° C for 10 min, hold at 4° C. PCR products were analyzed on a 0.8% agarose gel, excised and cleaned using the QIAquick gel extraction kit (Qiagen, 28115) and sent to Elim Biopharm for sequencing using the primers listed above.

#### Plasmids

“HIV-GFP” is full-length HIV encoding *GFP-IRES-nef* in the *nef* reading frame, also called “pNLENG-IRES”^62^ and “HIV-BFP” is the NL4-3 infectious clone^63^ encoding mTagBFP in the *nef* reading frame. Lentiviral vectors pLEG^64^, pLG^24^ and pFUGW^58^ have been previously described. The mutations for all F1 constructs and plasmids are listed in Supplementary Table 1. Cloning was done by various PCR cloning techniques as convenient and restriction cloning was used to exchange plasmid sequences as necessary. Sequences were confirmed by Sanger sequencing at Elim Biopharm.

### Transfection for viral stocks, lentiviral stocks or plasmid co-transfection experiments

For replication-competent HIV stocks, 293T cells were transfected with plasmid pNLENG1-IRES ^62^ or pNLENB-IRES. For F1 VLPs assembled in the absence of packaging plasmid, 293T cell were transfected with the indicated F1 construct. To functionally package F1 constructs or lentiviral vectors, the transfer plasmid, packaging plasmid pCMV-dR8.91^65^ and pseudotyping plasmid pCMV-VSVG (Addgene, # 8454) were mixed at a mass ratio of 4:10:5, respectively, except for F1 constructs prepared for primary cell transduction which were mixed at a 1:2:1 ratio. HIV:F1 plasmid co-transfection experiments were performed by mixing plasmids at the indicated molar ratios and equalizing the mass across all reactions with excess pUC19. To make the transfection mix, plasmids were diluted in unsupplemented DMEM (i.e., no serum or antibiotics added) to a concentration of 10 ng/mL total DNA and PEI was added to a concentration of 30 µg/mL in a volume ∼10% of the total volume in the transfection well or dish (e.g., 200 µl for a 6-well plate with 2 mL media). The transfection mix was vortexed vigorously, incubated for 10 minutes at room temperature, added to cells and incubated overnight. Transfection supernatant was replaced with fresh, supplemented DMEM the following day and supernatant was collected around 48 hours post transfection. Cell debris was removed by centrifugation and supernatants were filtered through a 0.45 µm filter. Concentrated virus was prepared by ultracentrifugation (Beckman Coulter Optima XE-90, rotor SW 32 Ti) at 20,000 rpm through a 6% iodixanol gradient (Sigma Aldrich, D1556-250mL) for 1.5 -2 hours.

### Titering HIV and lentiviral stocks

Replication-competent HIV (with fluorescent reporter) was titered by serial dilution on MT-4s and analyzed by flow cytometry on an BD LSR II or MACSQuant VYB flow cytometer at 2 days post infection. Lentiviral stocks with strong fluorescence expression (i.e., F1 constructs with Tat, and stocks made from pLEG or pFUGW) were titered as for HIV and reported as transduction units/mL. Lentiviral stocks with weak expression from the LTR (i.e., constructs without Tat) were titered by transducing MT-4s, superinfecting with full-length HIV at 2 or 3 days post transduction (to provide Tat in *trans*) and then measuring the percent of infected cells that were positive for lentiviral expression at 2dpi. All fluorescence data was analyzed in FlowJo™.

### Supernatant p24 concentration

The supernatant concentration of p24 was determined by the FLAQ method^56, 57^, essentially a sandwich ELISA performed on beads using a PE-linked antibody to enable quantification by flow cytometry. Briefly, serially diluted supernatant samples were incubated with KC57-PE p24 antibody (Beckman Coulter, 6604667) and polystyrene beads previously cross-linked to human anti-p24 Gag HIV-1 IIIB polyclonal antibodies (ImmunoDC LLC, 2503). After washing and fixing, bead fluorescence was analyzed by flow cytometry on a BD LSR II and fluorescence intensity was converted to concentration using a standard curve in FlowJo™.

### Transduction efficiency calculation

The efficiency of lentiviral stock transduction was measured as the transduction units (“titer”) per ng of p24 (i.e., capsid protein). Titers and p24 concentration were determined as described above.

### Construction of polyclonal cell lines

Polyclonal cell lines were created by transducing CEMs with lentivirus at low MOI (to ensure single integrations) and batch sorting GFP, BFP or mCherry+ cells (depending on construct used) with a BD FACSAria II cell sorter. Prior to sorting, populations transduced with F1 construct were treated with 5 µM Darunavir for 3-5 days to eliminate replication-competent recombinants. NB: Although GFP or BFP expression was very low for most F1 constructs due to the absence of Tat, dim expression was detectable on the BD FACSAria II. For doxycycline-inducible Tat cell lines, see below.

### T cell line infection and analysis

All cell line infections were performed by incubating cells with full-length replication-competent HIV-1 (pNL43 strain with a fluorescent reporter in the *nef* reading frame) at a density of 0.5 million cells/mL. The frequency of infection was monitored by fixing cells in 1% formaldehyde (Tousimis Research Corporation, 1008A), measuring fluorescent cells by flow cytometry on a BD LSR II and analyzing in FlowJo. For single cycle infection analyses, supernatants were collected at 3 dpi by pelleting cells, filtering supernatant through a .45 µM filter (Millex, SLHV033RS) and titering as described.

### Primary cell infection and analysis

T cells were purified from normal donors by negative selection using the RosetteSep Human CD4+ T Cell Enrichment Cocktails according to the manufacturer’s protocols (StemCell Technologies). Primary cells were activated and expanded using Human T-activator CD3/CD28 Dynabeads (Cat # 11161D) as directed by the manufacturer for 2 -3 days prior to infection. Cells were maintained in complete RPMI (10% FBS, 1% Pen/strep) with 30 U/mL rIL-2 (Peprotech, 200-02). To infect, activation beads were removed and one million cells were spinoculated in 100 μl total volume containing 1) 50 μl concentrated full length HIV-BFP, 2) 50 μl concentrated F1 (no Tat) lentiviral stock or 3) 50 μl each of concentrated full length HIV-BFP and F1(no Tat) at 1200g for 1.5-2 hours at room temperature. Following spinoculation, 100 μl media was added, cells were triturated and incubated overnight at 37 °C. The following day, cultures were expanded to 1 mL volumes with fresh media supplemented as described above. Fluorescent reporter expression was monitored by flow cytometry on an LSRII.

### Quantifying viral RNA and proviral DNA by allele-specific qPCR

Viral genomic RNA was extracted from filtered supernatants using the QIAamp Viral RNA Mini Kit (Qiagen, 52904) according to the manufacturer’s protocol. 2µl of a 1:1,000 dilution of MS2 RNA solution (Sigma-Aldrich, 10165948001) was spiked in as a normalization control. Extracted viral RNA was DNase treated with the TURBO DNA-free kit (Thermo Fisher Scientific, AM1907) according to routine DNase treatment protocol. Total cellular RNA was extracted using the Qiagen RNeasy Mini Kit (Qiagen, 74104) according to the manufacturer’s protocols including the on-column DNA digestion (RNase-free DNase I, Qiagen, 79254). RNA was reverse transcribed using either the QuantiTect Reverse Transcription Kit (Qiagen, 205311) according to the manufacturer’s protocols (including the gDNA wipeout step) or M-MuLV reverse transcriptase (NEB, M0253L) according to manufacturer’s standard protocol (using Murine RNase Inhibitor [NEB, M0314S]) including the recommended 25° C incubation for random primers, except that samples were denatured at 70° C and the enzyme was inactivated at 80° C for 5 min. For all cDNA synthesis, a no-RT enzyme control was included and reactions were primed with random hexamers. For integrated proviral, cellular DNA was extracted 2 days post infection using the NucleoSpin Tissue kit (740052.50) according to the manufacturer’s protocol. qPCR was performed using the Fast SYBR Green master mix (ThermoFisher Scientific, 4385612) according to manufacturer’s protocol using a 10 µl reaction volume and 200 nM final primer concentration on an Applied Biosystems 7500 fast real-time PCR system. Primers for total, full-length (i.e., HIV) and MS2 control RNA can be found in Supplementary Table 2. Because the efficiency for total and full-length primers was the same (97-101% efficiency, depending on experiment), the frequency of HIV RNA or DNA was determined by converting total and full-length Ct values to fold change (normalized to full-length only samples) and dividing full-length HIV fold change by total fold change. F1 RNA was assumed to be the remaining gRNA in the sample.

### Co-culture for F1 transmission

F1^cPPT^Δ-GFP or UBC-GFP transduced CEMs were infected with HIV-BFP and, at 2 days post infection, were co-cultured 1:20 with MT4 or CEM target cells expressing an mCherry reporter from the EF1A promoter. At various times post co-culture, cells were analyzed by flow cytometry on an LSRII flow cytometer with GFP/mCherry double positive cells indicating transmission of the F1^cPPT^Δ-GFP construct.

### Dox-inducible Tat cell line construction for measuring R_0_

To create a cell line that would transcriptionally activate F1 constructs lacking Tat expression (most of the constructs used in this work), we used the previously published doxycycline-inducible “Tat-FKBP” construct except that the Dendra reporter was exchanged for an mCherry reporter^66^. CEM cells were co-transduced with Tet-Tat-mCherry-FKBP and SFFV-rtTA (the transactivator required for doxycycline induction) lentivirus at high MOI. Cells were treated with 500 ng/mL dox for 24 hours prior to sorting the mCherry+ population and dox was washed out after sorting.

### Quantifying F1 transmission and calculating R_0_

On day 0, the F1^cPPT^Δ-GFP cell line (polycloncal, single integrations) or the UBC-GFP control cell line were infected with HIV-BFP at low MOI (to produce single integrations of HIV). At 3 days post infection, the frequency of BFP+ cells was measured by flow cytometry and 5,000 of these “producer” cells were co-cultured with 95,000 susceptible “target” cells. Target cells were the dox-inducible Tat-mCherry cell line described above treated with 500 ng/mL doxycycline 2 days prior to co-culture in order to activate transcription of any transmitted F1^cPPT^Δ constructs. At two days post co-culture, the frequency of HIV and F1^cPPT^Δ transmission was measured by flow cytometry for BFP/mCherry+ and GFP/mCherry+ cells, respectively.

To calculate R_0_, the following caveats were taken into account: 1) not all 5,000 “producer” cells were actually infected (only 12-19%), so the number of producers had to be adjusted to reflect this value. 2) uninfected “producer” cells (i.e., the original F1^cPPT^Δ cells that were not infected with HIV) are also technically susceptible to infection, so we assumed that these cells became infected at the same rate as the “target” cells in the second round of infection and added these cells into the calculation for the number of target cells in the culture. 3) Tat-mCherry expression from the dox-inducible promoter is not fully penetrant, even though the cell line was sorted for mCherry expression. Upon induction with dox, only about 50% of cells fall into the mCherry+ gate. Therefore, we used the mCherry gate to obtain the frequency of HIV or F1^cPPT^Δ secondary infections and then converted this to a number of secondary infections based on the known number of susceptible cells that were added to the culture.

R_0_ values were calculated as follows:

\# HIV-infected producer cells (i.e., primary infections) = frequency of BFP+ cells at 3 dpi x 5,000

\# HIV-infected target cells (i.e., secondary infections) = frequency of BFP+ cells in the mCherry gate x (95,000 + 5,000 - # HIV-infected producer cells)

\# F1-transduced target cells (i.e., secondary F1 “infections”) = frequency of GFP+ cells in the mCherry gate x (95,000 + 5,000 - # HIV-infected producer cells)

R_0_^HIV^ = secondary infections/primary infections

R_0_ ^F1cPPTΔ^ = secondary “infections”/primary infections

### Spinner flask cultures and F1 transmission

Following approximately 2 weeks of cell adaptation to constant agitation (see reactor protocol part A), cells were seeded in a 500 mL Wheaton bottle as follows: ∼13% F1^cPPT^Δ-GFP cell line and 87% CEMs expressing mCherry from the EF1A promoter (to mark them as target cells). Cells were infected with HIV-BFP at low MOI on day 0 and monitored for cell density, viability and fluorescence over time using a bio-rad TC20 and LSRII flow cytometer. Transmission of F1^cPPTΔ^-GFP was indicated by the double-positive mCherry/GFP+ population. The % infected cells was measured as the % BFP+ cells over time.

### Calculating percent interference

The percent interference was calculated by taking the change in titer relative to HIV only for each sample of interest (e.g., HIVonly titer - F1^cPPT^Δtiter) and dividing by the normalizing variant. For example, if comparing the interference observed with the F1^cPPT^Δ/ΔRT variant to that observed with F1^cPPT^Δ, divide both changes in titer by (HIVonly titer - F1^cPPT^Δ titer) so that F1^cPPT^Δ is set to 1, or 100% interference.

### Quantification of reverse transcriptase activity using the SG-PERT assay

We measured virion-associated reverse transcriptase activity using the published SG-PERT assay^55^ and the Fast SYBR Green master mix (ThermoFisher Scientific, 4385612). Briefly, virions were lysed and incubated with template RNA (MS2), primers for reverse transcription and qPCR mix. A one-step RT-qPCR was performed on an Applied Biosystems 7500 fast real-time PCR system, relying on virion reverse transcriptase proteins to generate cDNA.

### Western analysis

Cell-free virions were prepared by plasmid transfection and ultracentrifugation as described above. Virion samples were boiled in Laemmli buffer for 5 – 10 min and separated on a 4-12% SDS-PAGE gel. Proteins were transferred to a PVDF membrane using the Bio-Rad Trans-Blot SD Semi-Dry Transfer Cell (Catalog **#**1703940) at 10V for 25 minutes and probed as follows: membranes were blocked for 1 hour at room temperature with Li-Cor Odyssey blocking buffer in PBS (Catalog # 927-40000), incubated with a 1:2,000 dilution of Abcam anti-HIV1 p24 antibody [39./5.4A] (Catalog #9071) in blocking buffer supplemented with 0.2% Tween for one hour at room temperature or overnight at 4 degrees, washed with PBST 4 times for 5 minutes each, incubated with a 1:20,000 dilution of Li-Cor’s IRDye 800CW goat anti-mouse IgG secondary antibody (Catalog #926-32210) in blocking buffer supplemented with 0.2% Tween and 0.01% SDS, washed with PBST 4 times for 5 minutes each and analyzed on a Li-cor Odyssey imager using Image Studio acquisition software in the Western Analysis mode.

### HIV-1–humanized mouse challenge

6-week-old female NSG (NOD-*scid* IL2Rg^null^) mice were obtained from The Jackson Laboratory (JAX) and were injected via tail vein with 20x10^6^ human CD4 T cells in 100ul 0.5% human serum albumin in PBS. Prior to engraftment, TIP or GFP control transduced CD4 T cells were mixed with untransduced CD4 T cell so that 50% of the T cells were transduced. Three weeks later mice were tail vein injected with 15ng (p24) HIV Bal mixed 1:1 with PBS. Peripheral blood was obtained by retro-orbital bleeding, and human CD4 lymphocyte counts were enumerated using BD lysis buffer and BD TruCount tubes as previously described^67^, staining with mouse anti human CD45 PerCp Cy5.5 (BD Biosciences), and mouse anti human CD4 APC (Biolegend),

### Mouse serum RNA extraction

RNA was extracted from mouse serum using the BD TRI Reagent (Molecular Research Center, TB 126) according to manufacturer’s protocol for RNA extraction from serum. Before beginning the protocol, 150 µl DPBS was added to the ∼50 µl serum sample to bring the volume to ∼200 µl. Final resuspension of RNA was performed in 50 µl of nuclease-free water.

### ddPCR analysis of mouse RNA samples

Digital Droplet PCR was performed using the Bio-Rad QX100 droplet generator (1863002) and reader (1863003) according to the manufacturer’s protocol. RT-PCR reactions were carried out using the Bio-Rad One-Step RT-ddPCR Advanced Kit for Probes (BioRad, 1864021) following the manufacturer’s protocol with the following reaction conditions: 900 nM primers, 250 nM Taqman target probes and a 1X concentration of serum-extracted RNA. For thermocycling conditions, initial denaturation was 65°C for 5 min, and annealing and extension was performed at 56°C for 1 min, 5 sec. Data were acquired and analyzed with QuantaSoft Software. See Supplementary Table 3 for ddPCR primers and probes. All reactions were performed in technical duplicate and samples below the limit of detection (determined by negative controls) were not analyzed for F1 content. Viral loads were determined using primers and probes specific for full-length HIV. The frequency of HIV and F1 RNAs were determined with primer/probe sets specific for total RNA, full-length HIV RNA and F1 RNA.

### Statistical analysis

For data comparing frequency of infected cells, HIV titers, F1 transduction units or RT activity, p-values were obtained using an unpaired, two-tailed Student’s t test. For long-term culture infection trajectories, a paired, two-tailed t-test was used. For CD4+ T cell concentration in mouse experiments, intra- and inter-group comparisons were done using a Mann-Whitney test and viral loads were analyzed by one-way ANOVA and the Tukey’s test for multiple comparisons in Prism™.

## Ethics statement

Purified CD4 T lymphocytes were obtained by University of Pennsylvania Human Immunology Core/CFAR Immunology Core from de-identified healthy donors under IRB protocol 705906. All humanized mouse experiments were approved by the University of Pennsylvania’s Institutional Animal Care and Use Committee (Protocol 805606) and by ACURO (DARPA) and were carried out in accordance with recommendations in the Guide for the Care and Use of Laboratory Animals of the National Institutes of Health.

## Data availability

The data that support the findings of this study are available from the corresponding author upon reasonable request

## Biological materials availability

All unique biological materials are available from the corresponding author

## Code availability

Custom code is available upon request

## Acknowledgements

We thank Craig Mello, Douglas Richman, Jerome Zack, Keith Jerome, Melanie Ott, Warner Greene, and the Weinberger lab for discussions and suggestions. We thank Kathryn Claiborn for editing, Teresa Roberts for graphics support, and the Gladstone-UCSF CFAR flow cytometry core, supported by NIH P30 AI027763, S10 RR028962, and the James B. Pendleton Charitable Trust for technical support. The following reagents were obtained through the NIH AIDS Reagent Program, Division of AIDS, NIAID, NIH: MT-4 from Dr. Douglas Richman and CEM CD4+ cells from Dr. J.P. Jacobs^60^ (cat# 117). We also thank the UCSF Center for Advanced Technology access to ddPCR equipment and Eric Chow for advice in developing the ddPCR assay. This work was supported by the Bowes Distinguished Professorship, the Alfred P. Sloan Research Fellowship, the Pew Scholars in the Biomedical Sciences Program, the DARPA INTERCEPT program (D17AC00009), and the NIH Director’s New Innovator (OD006677) and Pioneer Award (OD17181) programs.

## Author contributions

E.J.T. and L.S.W. conceived and designed the study. E.J.T., J.L.R and L.S.W conceived and designed the mouse study. E.J.T., S.Y.J, C.T., J.G., Y.Z. and B.M. designed and performed the experiments, and curated the data. J.L.R. provided reagents. E.J.T. and L.S.W. wrote the paper.

## Competing interests

The authors declare that they have no competing interests

Supplementary information is included with this manuscript

