## Supplementary Methods for "Discovery and Engineering of a Therapeutic Interfering Particle (TIP): a combination self-renewing antiviral"

**TABLE OF CONTENTS**


1. **Numerical simulations of DIPs *in vitro***
   i. The model (equations, parameters, initial conditions) – **Page 2**
   ii. Model Result I: Serial-passage culturing of HIV-1 inhibits selection of spontaneously arising DIPs – **Page 5**
   iii. Sensitivity Analysis – **Page 6**
   iv. Model Result II: A continuous reactor culture (without target-cell addition) could allow for selection and amplification of spontaneously arising HIV-1 DIPs – **Page 7**
   v. Model Result III: Target-cell replenishment inhibits DIP expansion – **Page 8**
   vi. Modeling Takeaways: reactor design criteria – **Page 8**
2. **Numerical simulations of F1$^{cPPT}$ TIP *in vivo***
   a. F1$^{cPPT}$ TIPs could substantially lower HIV-1 set-point viral load – **Page 9**
3. **Reactor construction — detailed protocol**
   i. Materials and Reagents – **Page 11**
   ii. Equipment – **Page 11**
   iii. Procedures – **Page 12**
      a. Adapting suspension T-cell to chronic agitation – **Page 12**
      b. Preparation of parts – **Page 12**
      c. Assembly of peristaltic pump – **Page 13**
      d. Calibration of flow rate – **Page 15**
      e. Assembly of reactor vessel – **Page 15**
      f. Reactor maintenance – **Page 18**
      g. Reactor sampling – **Page 19**
      h. Reactor termination – **Page 20**
4. **Supplementary References – Page 21**

**NUMERICAL SIMULATIONS OF DIPS *IN VITRO***

### The model (equations, parameters, and initial conditions)

To predict how long it might take to select for (i.e., expand) an HIV-1 DIP that spontaneously evolves in culture (and understand why HIV-1 DIPs have not previously been reported), we adapted previously published dynamic models of DIPs for HIV-1[1-4] that are built upon the well-established 'basic model' of HIV in vivo dynamics[5,6]. Importantly, the DIP/TIP model introduces only two new parameters into the well-established basic model of HIV dynamics: an interference parameter ($\psi$) and a transmission parameter ($\rho$). All other parameters have been previously estimated [5,6] or were measured in our *in vitro* setting.

The DIP model consists of six coupled nonlinear ordinary differential equations (ODEs) describing the following state variables: target cells (T), HIV productively infected cells (I), HIV virion particles (V), DIP-transduced target cells ($T_t$), dually infected cells ($I_d$), and DIP virion particles ($V_t$).

We adapted the DIP/TIP model to describe *in vitro* cell-culture settings, and the equations are as follows (Eqs. 1):

$$\frac{d}{dt}T = \lambda + T\left(\frac{h-(T+T_t+I+I_D)}{h}\right) - d\,T - k\,T\,(V+V_t) - d_r\,T$$
$$\frac{d}{dt}I = k\,V\,T - \delta\,I - d_r\,I$$
$$\frac{d}{dt}V = N\,\delta\,I - \psi\,N\,\delta\,I_d - c\,V - d_r\,V$$
$$\frac{d}{dt}T_t = k\,V_t\,T\ +\ T_t\left(\frac{h-(T+T_t+I+I_D)}{h}\right) -\ k\,V\,T_T + P_r\,k\,V\,T - d\,T_t - d_r\,T_t$$
$$\frac{d}{dt}I_d = k\,V\,T_T - \delta\,I_d - d_r\,I_d$$
$$\frac{d}{dt}V_t = \rho\,\psi\,N\,\delta\,I_d - c\,V_t - d_r\,V_t$$

Briefly, as previously described[1-4], the equations describe T cells undergoing production ($\lambda$), division (represented by a canonical logistic-growth term with a 'carrying capacity' $h$), per capita death ($d$) and dilution ($d_r$) and infection by HIV virions (V) at probabilistic rate ($k$) which converts the cells to productively infected cells ($I$). I cells undergo virus-mediated cell death ($\delta$) and dilution ($d_r$) and generate $N$ virions V (i.e., the viral burst size). T cells are also infected by DIP virions ($V_t$) at rate $k$ equivalent to HIV infection rate that generates DIP-transduced cells $T_t$ which are functionally equivalent to T cells (divide and die at same rate). $T_t$ can be infected by HIV at rate $k$ (whereas I cells cannot be superinfected due to CD4 downregulation by HIV *nef*) and generate dually infected cell $I_d$. $T_t$ cells are also generated by spontaneous evolution of DIPs at a probabilistic rate $P_r$ dependent upon the level of wild-type virus infection. $I_d$ cells produce both $V$ and $V_t$ virions but the $V$ burst size is reduced by $\psi$ compared to I cells, whereas the $V_t$ burst size is altered by $\rho$ compared to the HIV burst size from these cells. In this *in vitro* setting all state variables undergo dilution at rate $d_r$.

The parameters, adapted to *in vitro* settings, are described in Table 1:

| PARAMETER | BIOLOGICAL INTERPRETATION | VALUE (UNITS) | JUSTIFICATION (references) |
|---|---|---|---|
| $\lambda$ | 'Basic model' parameter: T-cell (thymic) production rate | 30–50 cells/day<br>(0 for in vitro case) | References [5,6]<br>Set to zero for *in vitro* as no thymus or bone marrow replenishment, only cell division |
| $h$ | *In vitro* model adaptation: Carrying capacity within the logistic function: number of cells which population can sustain before division halts due to resource limitation | 2E6 (cells) | Measured *in vitro* for CEM and Jurkat cells in our culture settings as the maximum of cell concentration (per mL) that sustains efficient cell division; above this concentration division rate and cell viability begin to decrease |
| $d$ | 'Basic model' parameter: Death rate of uninfected $CD4^+$ T cells ($T$) | 0.02 ($day^{-1}$) | References [5,6] (but negligible contribution due to dilution factor $Dr$) |
| $d_r$ | *In vitro* model adaptation: Dilution rate due to *in vitro* passaging (i.e., media changes) | 0.2 ($day^{-1}$) | Controlled parameter |
| $k$ | 'Basic model' parameter: Infection rate of activated $CD4^+$ T cells per virion | 2.5E-8 $(virions \cdot day)^{-1}$ | References [5,6] |
| $\delta$ | 'Basic model' parameter: Death rate of HIV-1-infected cells ($I$) | 0.7 ($day^{-1}$) | References [5,6] (also measured in culture ref. [7]) |
| $N$ | 'Basic model' parameter: Burst size (no. of virions released from HIV-1-infected cell) ($I$) | 200 (virion/cell) | References [5,6] |
| $c$ | 'Basic model' parameter: Clearance (inactivation) rate of HIV-1 ($V$) and DIP ($V_T$) virions | 6.0 ($day^{-1}$) | *In vivo* measurements see refs. [5,6]; *in vitro* lentivirus inactivation $t_{1/2}$ measured to be ~4hrs (ref. [7]) |
| $\rho$ | DIP/TIP model parameter 1: Transmission parameter—fold difference in burst size of DIPs | Varied [from <1 to 2] | Varied to capture range of potential values (e.g., F1 DIP measured to have $\rho$ < 1, |

| | | | |
|---|---|---|---|
| | relative to HIV-1 from dually infected cells ($I_D$) | | whereas F1$^{cPPT}$ TIP engineered to have $\rho > 1$) |
| $\psi$ | DIP/TIP model parameter 2: Interference parameter—fractional therapeutic downregulation that DIPs exert on HIV-1 virion production from $I_D$ cells | Varied [0.1; 0.5; 0.9] | Varied to capture range of potential values (for comparison, $\psi$ for the F1$^{cPPT}\Delta$ TIP was measured to be 7/24 ≈ 0.3; see Fig. 2d, $R_0$ measurement) |
| $P_r$ | *In vitro* model adaptation: Probability of spontaneous DIP evolution/appearance in the viral population (Molecularly, the probability of a DIP being generated during HIV-1 replication) | $10^{-5}$ | Average HIV-1 error rate (ref. [8]) |

Initial conditions for most state variables were set to zero: I(t=0) = $T_t$(t=0) = $I_d$(t=0) = $V_t$(t=0) = 0. However, T(t=0) = $1.6x10^6$ to describe infection of a culture of T cells, and V(t=0) = $2x10^4$ to mimic inoculation of a culture at MOI ≈ 0.01.

Numerical solutions were obtained using custom code written in the Julia language (https://doi.org/10.1137/141000671) using a Rosenbrock (solution solver algorithm Rodas4 - https://openresearchsoftware.metajnl.com/articles/10.5334/jors.151/) with a relative and absolute tolerance at < $10^{-8}$. All the graphical representations were performed using Matplotlib in Python 3. Simulations were also checked in Mathematica™.

**<u>Modeling Result I</u>: Serial-passage culturing of HIV-1 inhibits selection (i.e., amplification) of spontaneously arising DIPs**

To mimic a serial-passage culture *in silico*, we used an iterative initial condition approach. The ODE system was numerical solved up until the time point of first passage ($t_p$) and then values were reset to initial conditions for the cell-related state variables (T[t], I[t], $T_t$[t], $I_d$[t]) to mimic supernatant-only transfer from well to well. However, we conserved the values from the previous iterations for viral particles state variables ($V_t$[t] and V[t]). We simulated a maximally conservative case with no *dilutions* of viral particles applied across serial passages (i.e., $d_r$ = 0 only in the $V$ and $V_t$ equations), thereby allowing the more permissive conditions for DIP emergence in the viral population. Initially we used $t_p$=10 but the DIP emergence was insensitive to reducing this parameter.

Representative simulations (using parameter estimates from Table I) are shown below.

For $\psi$=0.1 ; $\rho$=1:

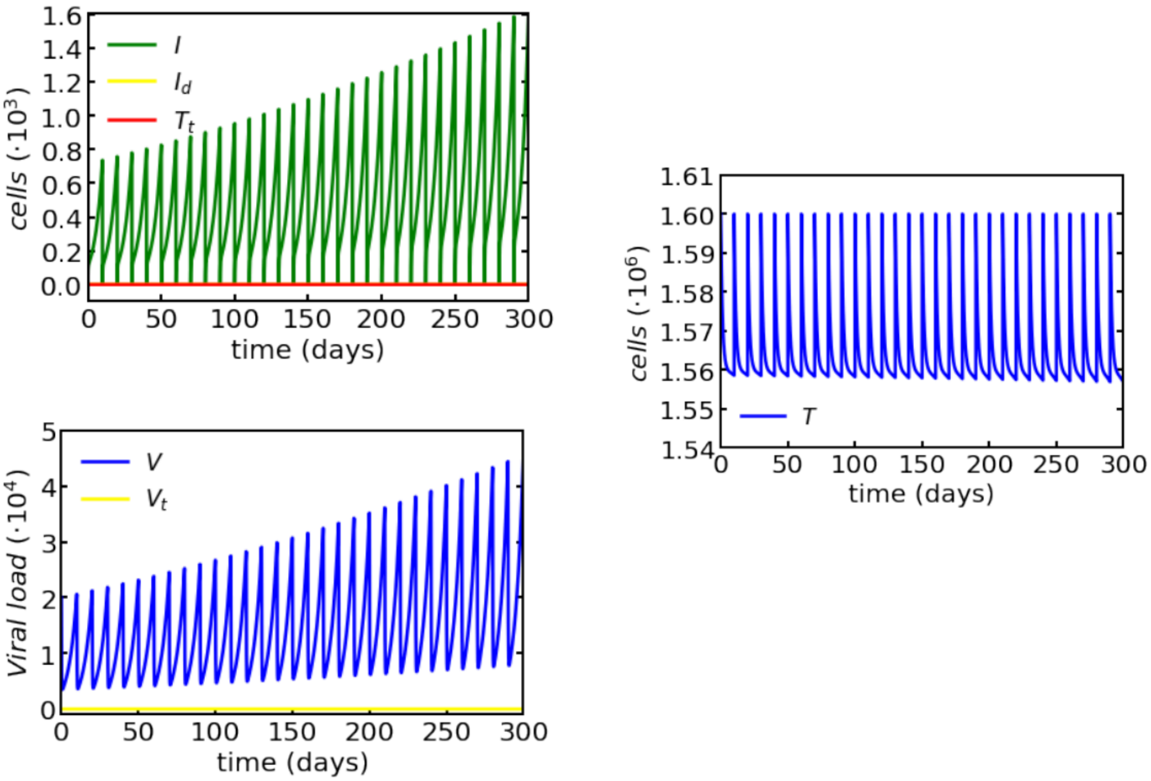


For $\psi$=0.5; $\rho$=1:

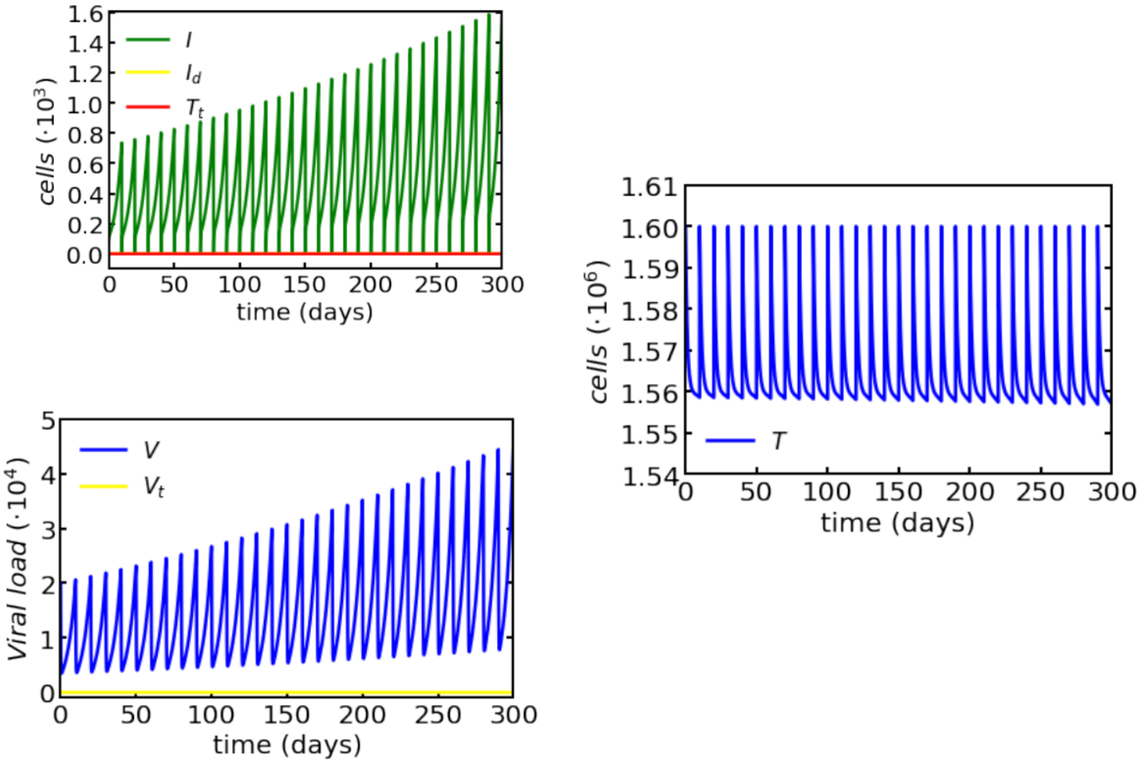


These simulation results—that a spontaneously arising DIP ($V_t$) would not expand—appeared to be general across parameter sets tested (i.e., parameter sensitivity analysis). For example, $V_T$ would not expand for changes in $\psi$ ($\psi$=0.9 gave similar results, not shown) or changes in $p_r$ ($p_r = 10^{-3}$ gave similar results). Below, we further explore this result.

**Sensitivity Analysis**:
We analyzed sensitivity of the model to several changes in parameters as well as several variations of the functional forms of the equations, including;

(i) accounting for virion removal—by adding virion-removal terms ($-\,k\,V\,T$ and $-\,k\,V_T\,T$) to the V and $V_t$ equations, respectively;

*(ii)* altering the functional form of the logistic growth term (e.g., by removal of the I and $I_d$ state variables from the summation term); and

*(iii)* adding the $d_r$ parameter to the V and $V_t$ equations (i.e., $\delta$ replaced by ($\delta$+$d_r$)).

None of these alterations of the functional forms generated a significant qualitative effect on the simulation results (i.e., there were small quantitative effects but no discernable qualitative effects on stability or kinetics). As detailed below, alteration of one key parameter, $\lambda$ (i.e., the fixed, constant rate of target-cell replenishment), did generate a significantly different and important effect (see Modeling result III section).

**<u>Modeling Result II</u>: A continuous-culture reactor (without target-cell replacement) could allow for selection and amplification of spontaneously arising HIV-1 DIPs**

We hypothesized that one reason for lack of DIP expansion, could be that replacement of excess target cells at each passage 'diluted out' the DIPs that spontaneously arose, disallowing them from gaining a foothold in the population. To test this *in silico* hypothesis we performed simulations where we did not reset the values at each $t = t_p$.

We note that these simulations are more conventional *in silico*, but they no longer mimic standard *in vitro* serial-passage cell cultures.

Using the parameter estimates in Table I for a range of $\psi$ values we obtained the following results:

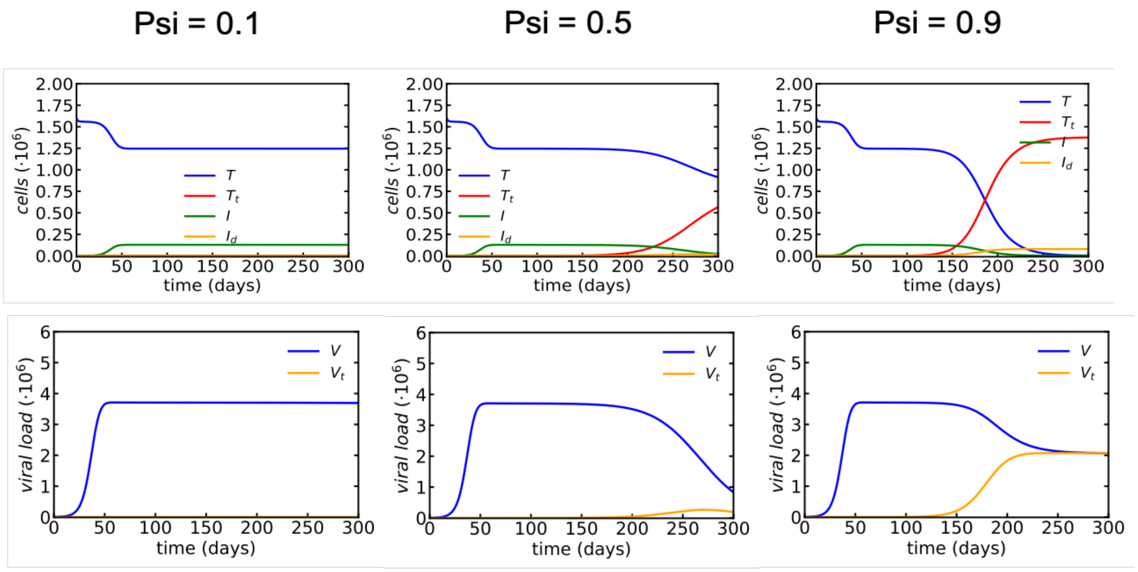


These results indicated that, under some parameter regimes, spontaneously arising DIPs could indeed expand in a culture setting were target cells were not replaced. However, it would take 100-200 days of continuous culture for the DIPs to expand. Under some changes of parameters, this time of arisal could be sped up slightly but we were unable to obtain DIP arisal times < 50 days in any of the models (not shown).

**<u>Modeling Result III</u>: Target-cell replacement inhibits DIP expansion**

To further test the hypothesis that replacement of excess target cells inhibited DIPs from expanding in the population, we set $\lambda$ (target cell production) to its non-zero literature-estimated value[5,6]. We obtained the following:

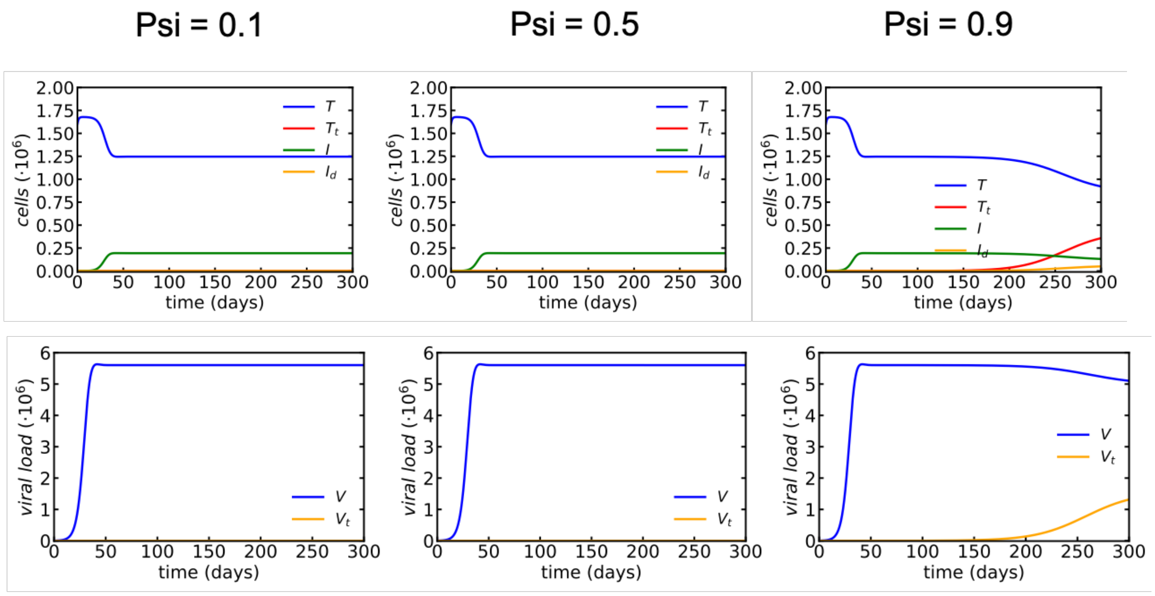


The simulations indicated that a constant addition of new target cells appears to delay DIP arisal time. The delay in DIP arisal was correlated with the value of $\lambda$ (not shown). Based on these data, we concluded that DIP arisal requires that target cells replenishment be provided only from division of existing cells.

**<u>Modeling Takeaways</u>: reactor design criteria**

The results above suggested to us that in order to isolate and expand a DIP for HIV-1 *in vitro* we would require:

1. A cell culture system that did not rely on conventional serial passage
2. A cell culture system where viral production could be maintained long term (i.e., 50-100 days)
3. A cell culture system that did not include target-cell replacement and where target cells replenishment is provided only by division of existing cells.

Based on these criteria, we examined a number of T cell lines (CEM, Jurkat, MT4) for their ability to divide and maintain an HIV-1 infection and found the CEM cells exhibited viral permissivity and a division rate that was the most likely to be compatible with supporting a long-term HIV-1 infection.

### NUMERICAL SIMULATIONS OF F1$^{cPPT}$ TIP *IN VIVO*

To predict the therapeutic effect of the F1$^{cPPT}$ TIP on HIV-1 set-point viremia in a patient we utilized equations from the most recent *in vivo* TIP model (ref [4]). The model is very similar to the system of ODEs presented above in Eqs. 1 but the *in vitro* parameters modified to reflect the *in vivo* situation and establish an HIV-1 set point. For example, the dilution rate was removed (i.e., $d_r = 0$), spontaneous evolution term was removed ($P_r$ set to 0), and, based on previous analyses[3,4], the model accounts for the likelihood that some $T_T$ could have more than one TIP integrated (i.e., $m$ TIP integrations) prior to being infected by HIV (i.e., $T_{T,m}$) and, and could divide prior to HIV infection (i.e., $I_{d,m}$). The functional form of the logistic growth term was also modified for the *in vivo* setting based on previous derivations[4]. The equation set retains the state variables as in Eqs. 1, with similar functional forms, and is as follows (Eqs. 2):

$$\frac{d}{dt}T = \lambda + T\,h_0\left(1 - \frac{T + \sum_m \mathrm{T}_{t,m}}{h_{\max}}\right) - d\,T - k\,T\,(V + V_T)$$
$$\frac{d}{dt}I = k\,V\,T - \delta\,I$$
$$\frac{d}{dt}V = N\,\delta\,I - \sum\nolimits_m \psi_m\,N\,\delta\,I_{d,m} - c\,V$$
$$\frac{d}{dt}T_{T,m} = k\,V_T\,T_{T,m-1} + T\,h_0\left(1 - \frac{T + \sum_m \mathrm{T}_{t,m}}{h_{\max}}\right) - k\,V\,T_{T,m} - d\,T_{T,m}$$
$$\frac{d}{dt}I_{d,m} = k\,V\,T_{T,m} - \delta\,I_{d,m} \qquad \left(for\ m = 1: T_{T,m-1} = T\right)$$
$$\frac{d}{dt}V_T = \sum\nolimits_m \rho_m\,\psi_m N\,\delta\,I_{d,m} - c\,V_T$$

Parameters are described in Table 1 except that values were varied within the ranges specified so that a 'pre-simulation' would match the physiological conditions that: (i) that only a small percentage of CD4$^+$ T cells are productively HIV infected during chronic set point infection; (ii) that acute infection and peak viremia transition to set-point viremia with ~30-60 days, and (iii) set point viremia stabilizes at ~100 (x10$^3$ copies/mL). The parameters set used to achieved these conditions is: {$\lambda$ = 30, $h_0$ = 3.3, $h_{max}$ = 3300, $d$ = 0.01, $k$ = 0.0005, $\delta$ = 0.7, $N$ = 100, $c$ =*30*} with initial conditions: {$T_{initial}$= $\lambda / d$ ; $V_{initial}$=0; $I_{initial}$=1}.

All TIP parameters and TIP-dependent state-variables were set to zero for the 'pre-simulation that was for the purpose of establishing the HIV-1 set point (i.e., $\rho_m$= $\psi_m$=0 and for all $m$ and initial values of $T_{T,m}$, $I_{d,m}$, and $V_T$, where all zero for all $m$).

The resulting set point viral load is $\bar{V}$=133.9 (note, this value is somewhat different than the typical $\bar{V}$= $(d/k)(R_0$-1$)$ where $R_0$=$\lambda kN/(dc)$=5, due to logistic-growth term).

Pre-simulation: establishment of HIV-1 set-point

For tractability, the simulation was started at t=–300 days so that TIP could be added at day 0 when simulation had reached stable steady state. The oscillations at early times are a known peculiarity of the simple HIV models that, for simplicity, neglect the adaptive CD8 immune response and other processes[6].

HIV viral load (no TIP):

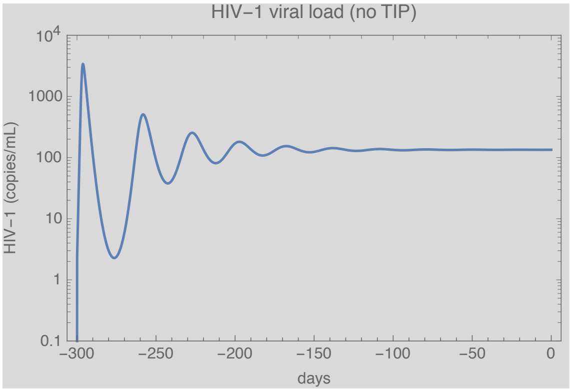


Effect of F1$^{cPPT}$ TIP on HIV-1 set-point viral load

Based on the results in Fig. 2D (main text), we used the measured $R_0$ values calculated the $\rho$ and $\psi$ values for the F1$^{cPPT}\Delta$ TIP. Given the HIV-1 $R_0$=24 being reduced to $R_0$=7 by F1$^{cPPT}$ TIP, we calculated $\psi_1=(7/24)$ and given the $R_0$=12 for the F1$^{cPPT}$ TIP, we calculated $\rho_1=(12/7)$. Based upon previous analysis[3,4], we used $\psi_m=\psi_1$ *and* $\rho_m=m\rho_1$. *[m* was limited to a ceiling of $10^1$ for computational tractability].

The initial conditions were taken from the pre-simulation above and a bolus of TIP was introduced (either via $T_T[0]$=1000 or $V_T[0]=10^6$).

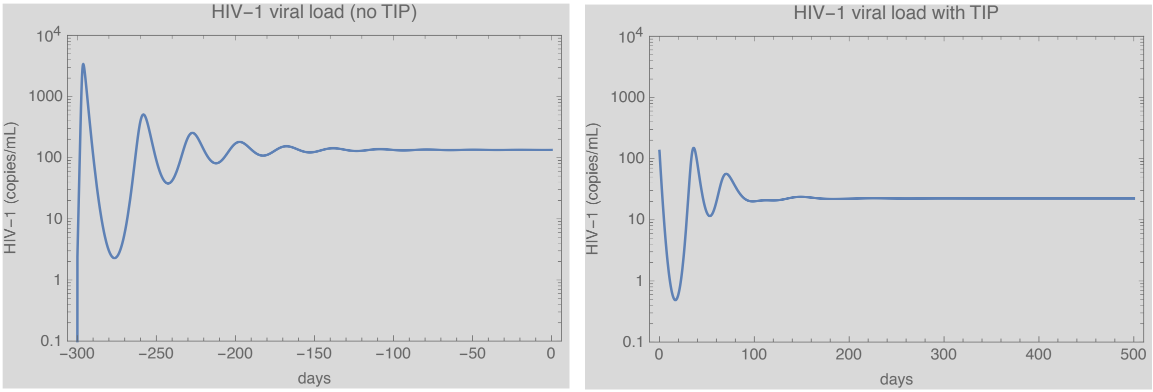


In agreement with previous predictions[1-4], the simulations predict that introduction of **F1$^{cPPT}$ TIP results in a ~1-Log reduction in HIV-1 set point** ($\bar{V}$=134 copies/mL → $\bar{V}$= 29.2 copies/mL).

Epidemiological models project[1-4] that 1 Log set-point reduction could substantially lower population-level prevalence of HIV-1.

**REACTOR CONSTRUCTION PROTOCOL**

### Materials and reagents

1. CEM CD4+ Cells (NIH AIDS Reagent Program, 110135)
2. RPMI 1640 with L-Glutamine (Fisher Scientific, MT10040CV)
3. Fetal Bovine Serum, FBS (Fisher Scientific, MT35011CV)
4. Penicillin-Streptomycin Solution, Pen/Strep (Fisher Scientific, MT30002CI)
5. Vented screw caps, two sizes (Fisher Scientific, 07-201-306 and 08-774-205)
6. Peristaltic Pump Tubing, red-red (Elemental Scientific, MPP-114-F-PVC)
7. PFA tubing 1/16" OD x .030" ID x 50ft (IDEX Health & Science, 1514L)
8. Luer adapter 1/4-28 female to female luer (IDEX Health & Science, P-658)
9. Flangeless male nut, 1/4-28 flat-bottom, 1/16" OD (red, white, and green) (IDEX Health & Science, P-202, P-203, P-205)
10. Flangeless ferrule, 1/4-28 flat-bottom, for 1/16" OD (IDEX Health & Science, P-200)
11. Peristaltic pump tubing adapter (IDEX Health & Science, P-757)
12. MicroClave port male adapter plug (Med Supply, HOS-12568-01)
13. Wheaton Celstir Spinner Flasks, 125 mL and 500 mL (Wheaton, 356876 and 356882)
14. Omnifit "Q" series bottle cap (Fisher Scientific, 00932Q3V)
15. Aluminum foil
16. Labeling tape
17. Kimwipes (Kimtech Science, 34133)
18. Ethanol, 190 proof (Koptec, V1101)
19. 20% Formaldehyde (Tousimis Research Corporation, 1008A)
20. 0.45 µm sterile syringe filters (Millex, SLHV033RS)
21. 10mL disposable sterile syringes with Luer-Lok™ tips (Fisher Scientific, 14-823-16E)
22. 

### Equipment

1. Cell counter *(Bio-Rad, TC20 Automated cell counter)*

2. Flow cytometer (LSR II)
3. 4-position slow-speed stir plate *(Fisher Scientific, 11-495-03)*
4. Micro peristaltic pump with manual speed control, 8-channel *(Elemental Scientific)*
5. Capillary polymer tubing cutter *(IDEX Health & Science, A-350)*
6. Timer
7. Awl
8. Scissors

**PROCEDURES**

A. Adapting suspension T cells to chronic agitation
   1. Culture CEM cells (*NIH AIDS Reagent Program, 110135*) in complete RPMI (10% FBS, 1% Pen/Strep) in an autoclaved 500mL stir flask (*Wheaton 356882*) with ventilated screw caps (*Fisher Scientific, 07-201-306)*. Cells should not exceed a density of 2 million/mL.
   2. Place the stir flask on the slow-speed stir plate (*Fisher Scientific, 11-495-03*) inside of a 37°C incubator with 5% $CO_2$ and stir at 90 – 120 rpm.
   3. Monitor cell concentration and viability, diluting to ~0.5 million/m when they reach confluency, for 1-2 weeks to ensure that the cells adapt to the chronic agitation. Adapted cells should double approximately every 24 hours with >80% viability.

B. Preparation of parts:
   1. Autoclave the following parts to construct a single virostat flask (we often run four at once):
      - 125 mL stir flask (*Wheaton, 356876)*
      - PFA tubing 1/16" OD x .030" ID x 50ft *(IDEX Health & Science, 1514L)*
      - 4 peristaltic tubing adapter sets *(IDEX Health & Science, P-757)*
      - 4 flangeless male nuts, 1/4-28 flat-bottom, 1/16" OD (red, white, and green) *(IDEX Health & Science, P-202, P-203, P-205)*

- 4 flangeless ferrules, 1/4-28 flat-bottom, for 1/16" OD *(IDEX Health & Science, P-200)*
- 1 luer adapter 1/4-28 female to female luer *(IDEX Health & Science, P-658)*
- 1 Kinesis™ Omnifit™ "Q" series bottle cap (*Fisher Scientific*, 00932Q3V)
- 2 caps salvaged from used media bottles (one cap for media bottle and one for waste bottle) with holes (use a Dremel or awl), one hole per virostat flask. The holes need to be small enough to tight-fit the PFA tubing with 1/16" outer diameter.

2. Other parts and tools (not autoclaved):
   - MicroClave port male adapter plug *(Medex Supply, HOS-12568-01)*
   - 2 PVC peristaltic pump tubes, red-red *(Elemental Scientific, MPP-114-F-PVC)*
   - 1 Corning vented screw cap *(Fisher Scientific, 08-774-205)*
   - Micro peristaltic pump with manual speed control, 8-channel *(Elemental Scientific)*
   - biohazard stickers
   - Tin foil
   - Tape for labeling flasks, sample port lines and taping down flasks
   - Capillary polymer tubing cutter *(IDEX Health & Science, A-350)*
   - Pliers
   - Scissors

C. Assembly of peristaltic pump parts:

1. Place all autoclaved and sterile components inside of a biosafety cabinet in an approved BSL-3 facility.
2. Assemble peristaltic pump tubing adapters on one end of the peristaltic tubing. Use pliers to securely fit the tubing end onto the adapter and tighten adapter. Repeat for other end of tubing (see examples below).

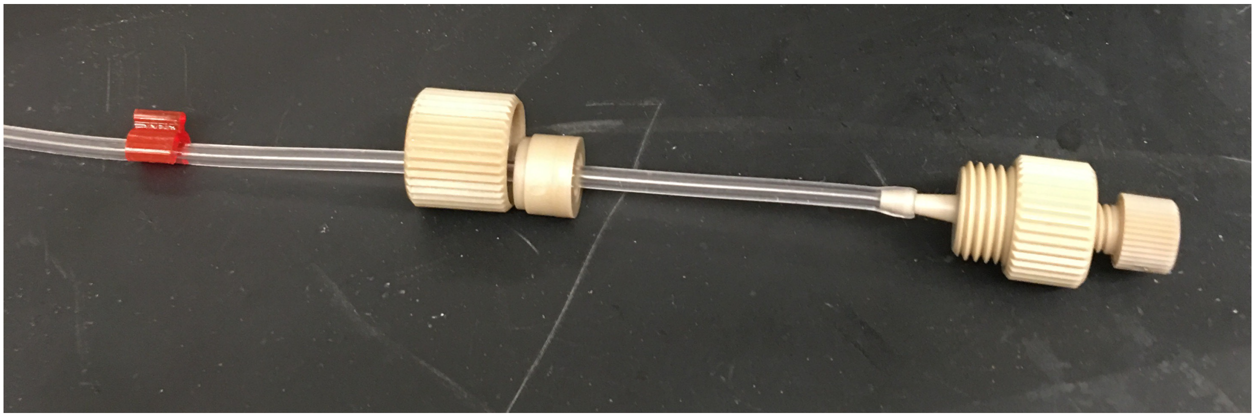

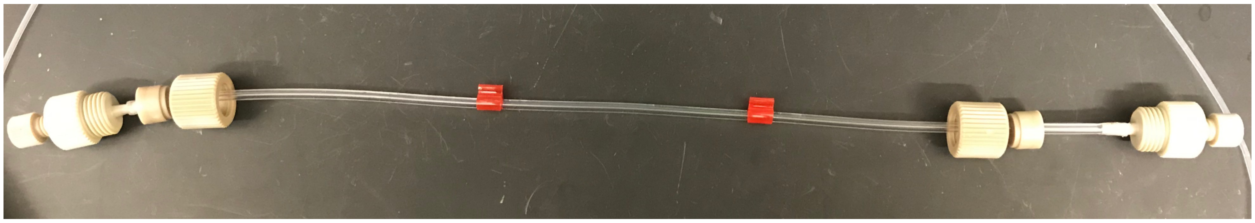

3. Repeat step 2 for a second set of tubing.
4. Cut PFA tubing into 4 lengths, long enough to reach from peristaltic pump tubing to stir flask or media/waste bottles. Create flush tubing ends using the tube cutter.
5. Tightly connect a length of the PFA tubing onto each end of the peristaltic pump tubing using the adapters (see image to right). Thread the PFA tubing through the free end of the adapter and then the ferrule and screw into place.

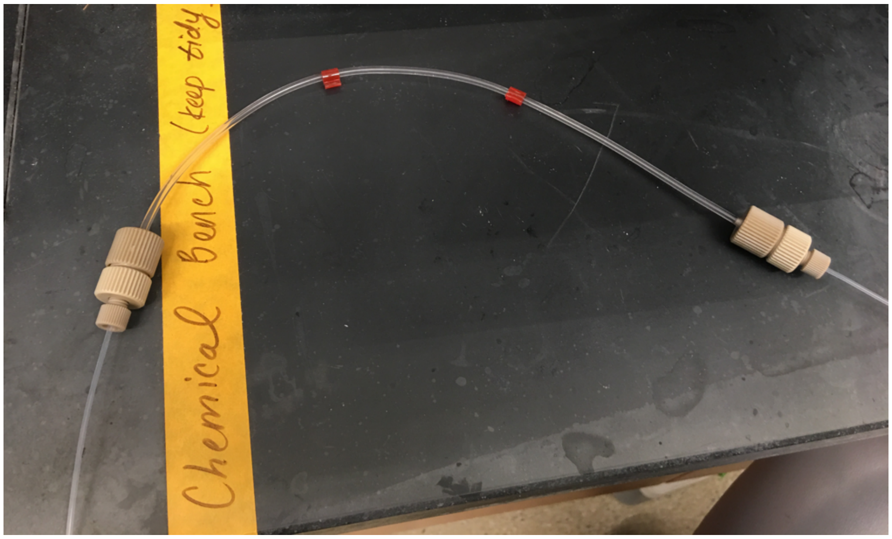


**Note**: at this point you should have two sets of liquid lines per flask (one to pump in fresh media and one to pump out waste).

6. Add assembled lines to peristaltic pump (see image at right.)

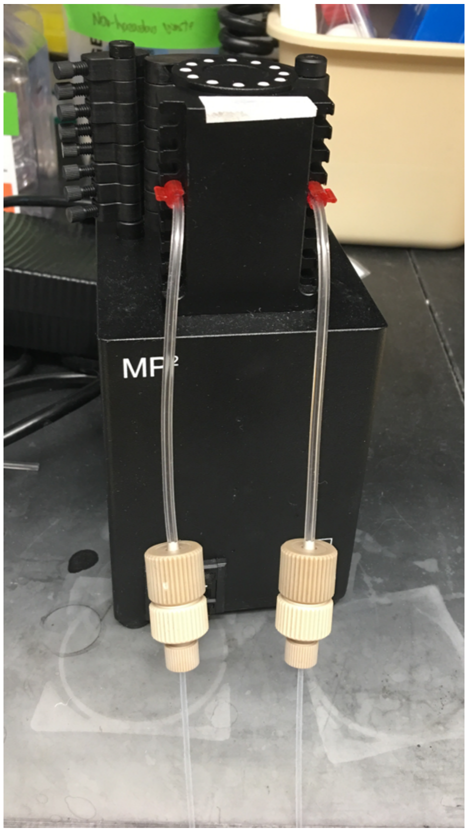

D. Calibrating flow rate of peristaltic pump:
1. Add inlet lines to clean water vessel
2. Add outlet lines to individual collection tubes
3. Turn pump on
4. Confirm that all fluid lines take up and dispense water
5. Run fluid through lines for 30 min at about 1 mL/min
6. Check that all fluid lines are within 5% error (by volume or weight)
7. Adjust nobs (tighten to speed up, loosen to slow down) to recalibrate as necessary
8. Run ethanol through lines to clean
9. Run clean water to remove ethanol
10. Cover ends of liquid lines with clean aluminum foil to prevent touch contamination

E. Assembly of a reactor flask (repeat for each desired flask):
1. Screw a "Q" series screw cap on one of the two arms of a 125 mL stir flask. Screw on a new vented screw cap on the other arm. Label flask with contents and biohazard sticker
2. Add the liquid delivery line to the Q series screw cap by threading the end of the liquid delivery line through a male nut, a ferrule, and then the port in cap (see image below). Screw the nut into the port. Be sure liquid will flow *into* the flask

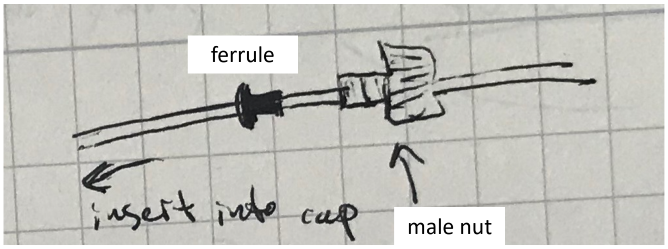

**Note**: The end of the media delivery line should ***not*** be submerged in the culture (once cells are added) in order to avoid contamination

3. Add the waste removal line by repeating the above step except be sure the liquid will flow *out* of the flask.

    **Note**: The end of the removal line should be submerged in the culture (once cells are added) to remove culture.

4. Add a sampling line (for taking timepoints, etc) by cutting a length of PFA tubing, creating flat ends with the tube cutter and threading through the Q series cap port as described in step 2.
5. Connect the other end of the sampling line to a microclave port by threading it through a male nut and ferrule, then screw into a female-to-female connector and screw onto the microclave port (see drawing below). Cover in aluminum foil to keep sterile. These MicroClave outlets will allow samples to be collected without moving the stir flasks.

    **Note**: can keep screw loose in order to move sampling line into and out of culture at willl

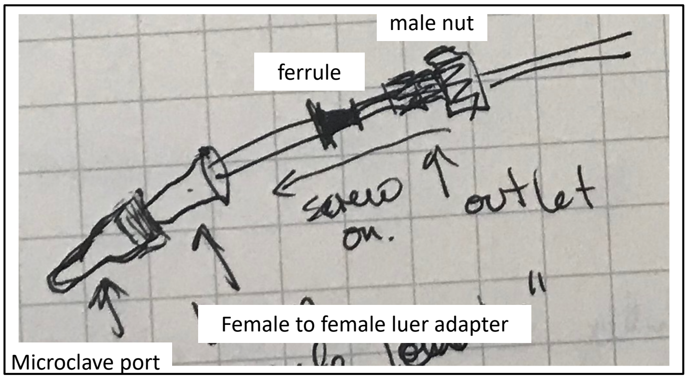

6. Add free ends of media delivery lines and waste lines to media and waste bottles, respectively, by spraying tubing ends with ethanol, drying with a clean Kimwipe and threading free tubing ends through pre-made holes in bottle caps. Be sure media lines are submerged to bottom of media bottle.

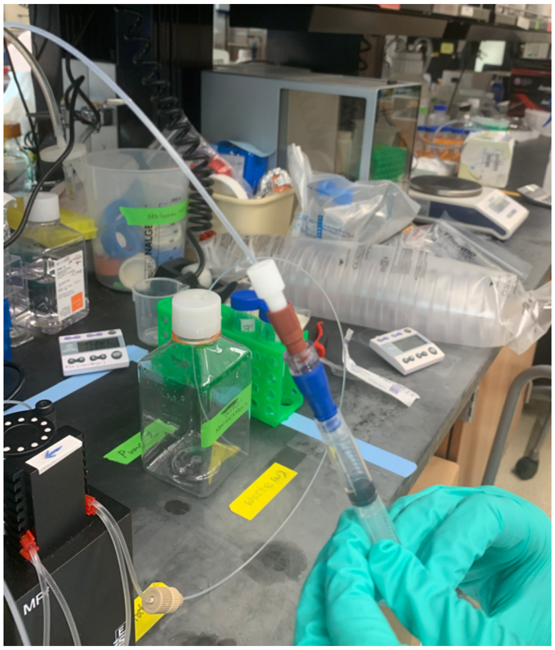

7. Check that all lines run in expected direction and test sampling ports.
8. Run media through lines for 10-30 min to clear remaining water.
9. Transfer cells from Part A to the flask, seeding at a density of ~1x$10^6$ cells/ml in a total volume of 80- 125 mL. Mark the solution level on the surface of the flask as well as media and waste bottles.
10. Collect all components (flask, pump, media and waste) in a large tub and transfer entire apparatus to a 37˚C cell culture incubator inside of an approved BSL-3 facility.
11. Place the stir flask onto the stir plate and tape it down to prevent tipping over. Turn the stir plate on to operate at 90rpm – 120rpm.

12. Calculate the pump speed from the calibration results and program a timer to operate the pump periodically to achieve a .25 dilution rate/day. For example, for a 120 mL culture and at a 1 mL/min flow rate should, the pump should operate for 5 minutes every 4 hours.

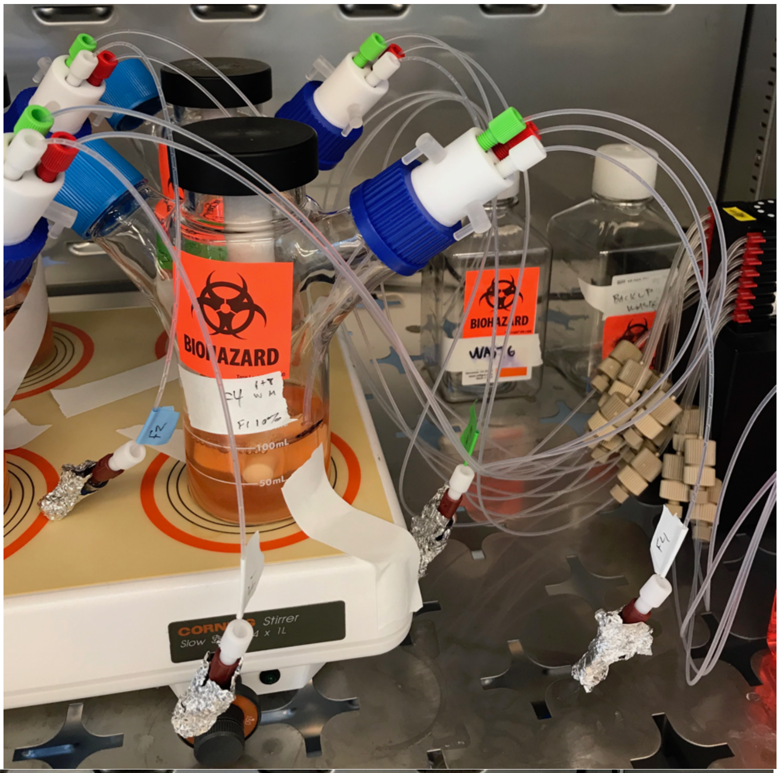


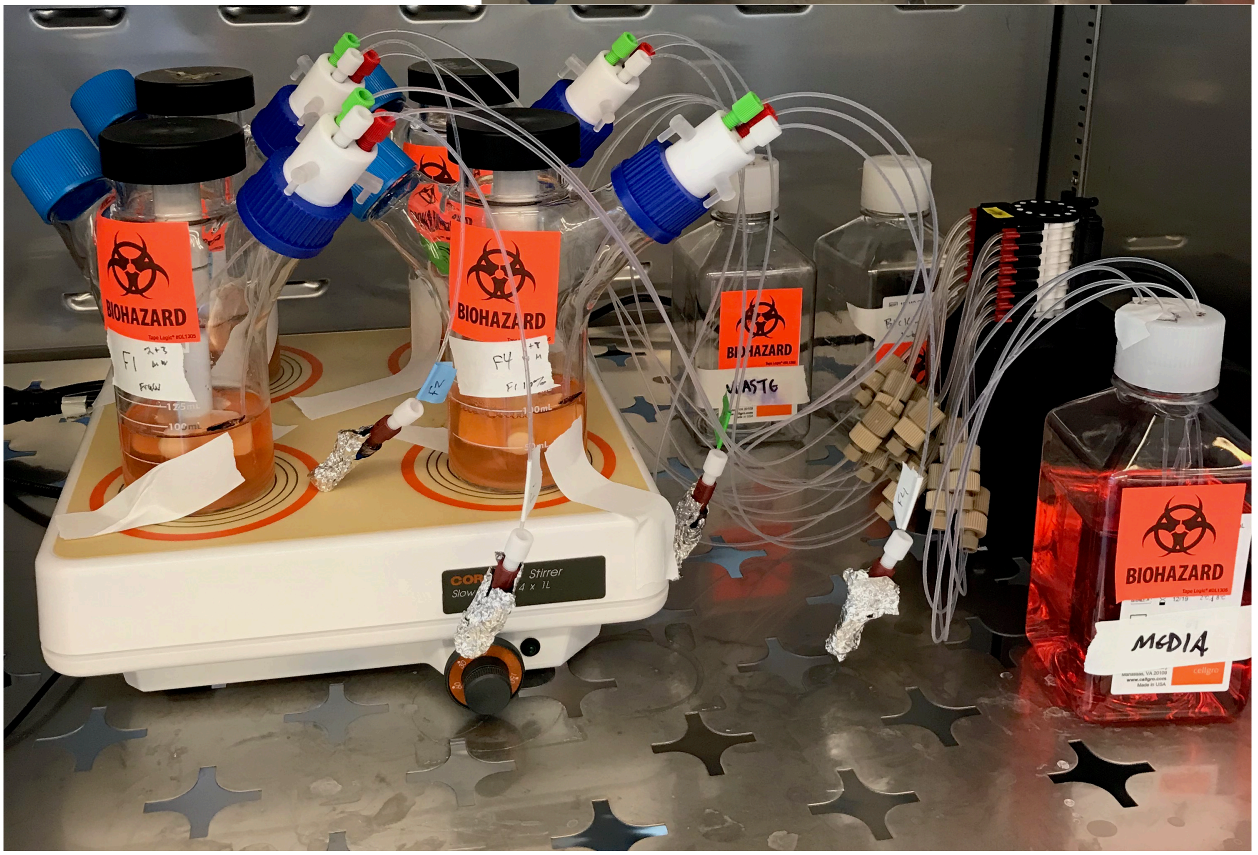


F. Reactor maintenance:

1. Check level in media bottle; replace with new media bottle if necessary
2. Check culture volumes in flasks; add back media to designated volume line if necessary (use the microclave port to introduce fresh media with a syringe)
3. Measure waste volume:
   - In incubator, carefully unscrew waste bottle cap and screw onto backup waste bottle
   - Use backup bottle cap (no holes) to close waste and remove to biosafety cabinet
   - Measure waste volume and dispose waste in bleach bucket

G. Reactor sampling:

1. Acquire sample from each virostat flask:
   - Prepare collection equipment: for each flask, prepare a 10 mL collection syringe, pull plunger to 3 mL line and rest into 50 mL falcon collection tube
   - Bring collection syringe to incubator, remove foil from end of one sample port and screw on 10 mL syringe
   - Remove 3 mL of flask contents, keeping syringe vertical (so culture collects at plunger interface with air gap at top)
   - While still screwed in, push syringe plunger so that air is introduced into the sample port line (to clear liquid from the line, reducing probability of a biohazardous spill/drip)
   - Unscrew syringe, place in 50 mL falcon and cover the end of the sample port with foil
   - Repeat for each flask
   - Bring samples back to biosafety cabinet and add sample from syringe to 50 mL conical for downstream processing
2. Fix cells and analyze fluorescence by flow cytometry.
3. Determine cell count and viability (we use a Bio-rad TC20)
4. Harvest culture supernatant by pelleting cells, filtering supernatant through a 0.45 μm filter and freezing at -80.
5. Freeze cell pellets for future gDNA extraction and analysis

H. Reactor termination

1. Fill up the media bottle with 20% bleach. Turn on the peristaltic pump to operate at its full speed. Let the system bleach out completely (overnight).
2. When the entire system is bleached, run the distilled water for 1 day.
3. Remove all of flasks from incubator and spray 70% EtOH thoroughly.
4. Empty out and rinse all the flasks and bottles.
5. Spray 70% EtOH to the rest of equipment in the incubator.
6. Thoroughly wipe down the incubator interior.
7. Autoclave all of flasks, tubing and parts to store.
